# The anterior cingulate cortex drives lateralized age-dependent modulation of claustrum circuits

**DOI:** 10.1101/2025.09.09.675132

**Authors:** Tarek Shaker, Mohammed Almokdad, Lotte JAM Razenberg, Mahesh M Karnani, Jesse Jackson

**Affiliations:** Department of Physiology, University of Alberta.; Neuroscience and Mental Health Institute, University of Alberta, Edmonton, Alberta, Canada, T6G2H7; Department of Physiology and Biophysics, State University of New York at Buffalo, Buffalo, NY 14203; Department of Integrative Neurophysiology, Center for Neurogenomics and Cognitive Research, Vrije Universiteit Amsterdam, Amsterdam Neuroscience, 1081 HZ Amsterdam, the Netherlands; Institute for Neuroscience and Cardiovascular Research, The University of Edinburgh, Edinburgh EH8 9XD, Scotland, The United Kingdom

**Keywords:** Development, GABA, Claustrum, Top-down, Prefrontal cortex, Interneurons

## Abstract

The anterior cingulate cortex (ACC) sends top-down inputs to the claustrum during sensory, motor, and cognitive processing. This ACC input is thought to drive the activation of claustrum neurons which in turn project back to the cortex to help orchestrate cortical networks during demanding cognitive states such as attention. However, the circuit mechanisms underlying ACC-claustrum signaling are not fully understood. Using in vivo single neuron recordings in mice, we show that ACC neuron activation drives a lateralized modulation of claustrum excitability that changes as a function of postnatal age. In adulthood, ACC activation evoked feed-forward inhibition of ipsilateral excitatory claustrum neurons and activation of contralateral excitatory claustrum neurons. Chemogenetic manipulation in adult mice revealed that ipsilateral claustrum inhibition by the ACC was due to feed-forward activation of claustrum parvalbumin inhibitory neurons. However, in neonatal mice, which lack mature parvalbumin interneurons, ACC inputs evoked claustrum excitation. In juvenile mice, the developmental switch from ACC-evoked claustrum excitation to inhibition occurred in parallel with the maturation of claustrum parvalbumin interneurons, thus corroborating the chemogenetic findings. Therefore, this work provides a novel mechanism of cortical control over claustrum activity that is refined during early postnatal life.

## Introduction

The anterior cingulate cortex (ACC) is a high-order cortical region involved in a myriad of different functions including attention allocation^1,2^, sensory processing^3^, and predictive coding^4–6^. Pathological conditions such as chronic pain and stress are associated with dysfunctional ACC neural activity^7–9^. ACC neurons send top-down inputs to several cortical and subcortical brain regions including the claustrum^10–14^. The claustrum and ACC communicate bidirectionally, using reciprocally coupled loops that are thought to be involved in motor planning^15^, stress regulation^16,17^, cognitive control^18,19^, sleep^20–25^, memory^22,26–29^, and pain^30–33^. However, the physiological circuit mechanisms underlying top-down ACC-claustrum communication remain incompletely known. Understanding this circuit will inform its role in an array of functions and how pathological states could be treated through circuit specific interventions.

Previous anatomical and physiological experiments have shown that excitatory neurons in the ACC send bilateral projections to the claustrum^11,30,34–37^ which synapse on both excitatory and inhibitory neurons^10,12,38–40^. Optogenetic activation of ACC axons in vitro can drive supralinear responses from claustrum neurons – where claustrum neurons fire at frequencies exceeding the frequency of ACC stimulation^12,40–42^. Claustrum neurons send projections back to the cortex of the ipsilateral hemisphere, densely innervating the ACC and most other frontal, midline, and temporal cortical regions^10,13,36,43–48^. These circuit properties have informed theories of claustrum function. For example, it has been hypothesized that claustrum circuits amplify and/or integrate multisensory information from different cortical regions and relay the message across cortical regions^18,19^. Previous work on ACC-claustrum connectivity has been done in vitro; therefore, it is necessary to understand how this circuit operates in vivo to determine the physiological principles underlying ACC-claustrum processing.

To address this knowledge gap, we performed in vivo extracellular recordings in mice to determine how claustrum neurons are modulated by the ACC, across different postnatal ages. The functional effect of ACC on claustral circuits was lateralized, and, surprisingly, ipsilateral ACC inputs were attenuated rather than amplified by claustrum circuits. In adult mice, ACC inputs drove the suppression of claustrum excitatory cells in the ipsilateral hemisphere, and activation in the contralateral claustrum. This feed-forward inhibition in the ipsilateral claustrum was mediated by parvalbumin-expressing (PV) interneurons and was absent in neonatal mice prior to claustrum PV cell maturation. These results help provide clarity regarding what role claustrum circuits play in claustrum-cortical loops and show an age-dependent hemisphere specific modulation of claustrum activity which may be used to support top-down communication from the ACC to other cortical regions.

## Results

To study how claustrum neurons respond to ACC inputs in vivo, we injected the ACC of wildtype mice with channelrhodopsin2 (ChR2) restricted to excitatory cortical neurons (AAV1-CAMKII-ChR2-eYFP) (**Figure 1A-D**). Following 14-21 days of expression, we performed in vivo extracellular recordings from either the ipsilateral or contralateral claustrum during ACC stimulation in mice under light isoflurane anesthesia (**Figure 1E-F**, see **Methods and Materials**). Great care was taken to ensure our electrodes were located within the claustrum using post-experimental histological verification of all electrode locations. To do this, we defined the claustrum anatomically as previously done in vitro using the dense fluorescence of ACC-ChR2 axons that demarcate the claustrum from surrounding cortical and striatal regions^41,49^. We also used the cortical marker Tle4 which is absent from classic claustrum neurons^50,51^ and nearly absent from the anatomical region we defined as the claustrum (**Figure 1C-D**). Electrodes were coated in Dil or DiD, to confirm that the recording location of every neuron was within the region defined by ACC-ChR2 axons and lacking Tle4. (**Figure 1G, Methods and Materials**). Cells recorded that could not be spatially verified were not used. In total, 46 mice were used and analyzed in this way to obtain a total of 149 single neurons in the claustrum and 29 cells “off target” in layer 5 of the insula (**Methods and Materials**).

**Figure 1:**
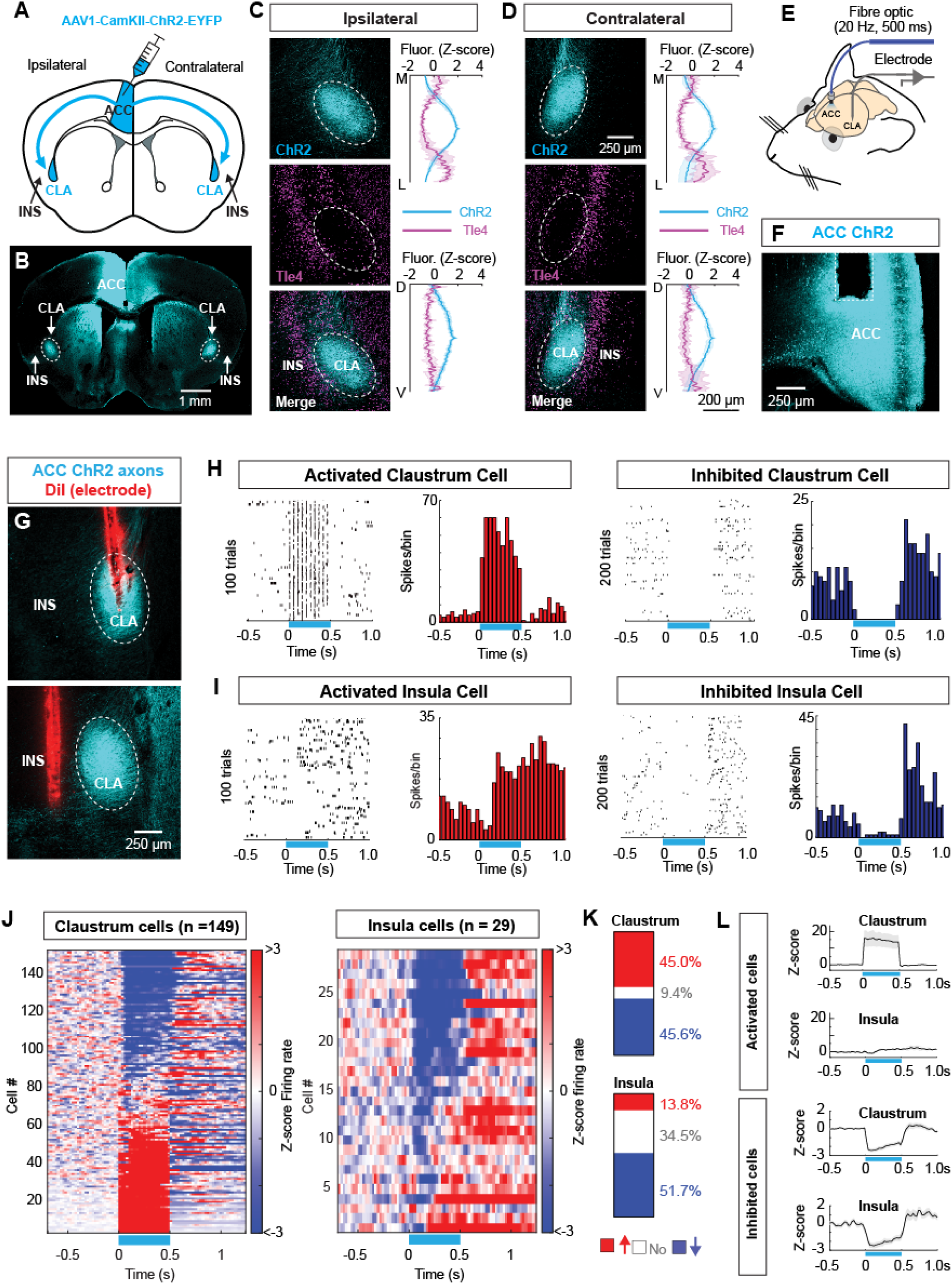
The anterior cingulate cortex drives both strong excitation and strong inhibition in claustrum neurons. A: Schematic of the experimental setup, where virus is injected into the anterior cingulate cortex (ACC). B: Representative image showing the expression of ChR2-eYFP in the injection site (ACC) and axons throughout the striatum and claustrum. Claustrum and insula regions are highlighted. C-D: Representative image of ACC axons innervating the ipsilateral (C) and contralateral claustrum (D). Tle4 is a cortical marker not expressed in claustrum neurons, as used as an additional spatial marker for the presence of the claustrum. Fluorescence intensity plots of ACC axons (ChR2) and Tle4 are shown in the medial (M)-lateral(L) and dorsal(D)-ventral(V) axes. E: Schematic of the setup used to record claustrum neurons in vivo while stimulating ACC cell bodies. F: Representative image showing the placement of the optic fiber cannula placement into the ACC. G: Example images showing the location of the recording electrode labelled with Dil in the claustrum (top) and insula (bottom). H-I: Example peri-stimulus time histogram. Spike times across stimulation trials are plotted in black, and the summed spike count potted in red (activated cells) or blue (inhibited cells). Example claustrum neurons that were excited (top left) or inhibited (top right), and insula cells that were excited (bottom left) or inhibited (bottom right). 100-200 trials are shown but, in many cases, >500 trials were performed. The horizontal blue line indicates stimulation period. J: Heatmaps showing the complete data set from adult mice, where the response of all neurons verified as being in the claustrum (left) or insula (right). Data were collected in 46 mice (25 female, 21 male). K: The proportion of claustrum and insula neurons that were activated, inhibited, or unchanged. L: The mean z-scored firing rate for all claustrum and insula cells that were activated (top) or inhibited (bottom), noting the activated insula cells are less strongly modulated than claustrum.

First, we compared the response of neurons located in the claustrum and insula, pooled across cell types and hemispheres in adult mice. In the claustrum, a similar percentage of neurons were activated (45%) or suppressed (46%) by the 20Hz ACC stimulation (**Figure 1H-K**). This contrasts with in vitro results where nearly all claustrum neurons are activated by ACC axonal stimulation, often spiking in excess of 100Hz ^40,41,49^. High firing rate responses were never observed here in vivo, and only 11/149 neurons (7%) in the claustrum data set exceed the 20Hz stimulation frequency, showing ACC inputs are attenuated by the claustrum. Control mice that received the same light pulses with no functional opsin did not exhibit meaningful changes in claustrum cell firing (**Figure S1**). Although the insula is not densely innervated by the ACC in mice^34^, cells recorded in the insula were predominantly suppressed (52%) and activated responses (14%) were weak relative to excitatory claustrum responses (**Figure 1I-K, Figure S2**). Overall, these data show that the firing of claustrum neurons can be either increased or decreased by ACC stimulation and that insular cortex is predominantly inhibited by ACC inputs (**Figure 1L**). The bimodal modulation (excitation and inhibition) of claustrum activity was not shared by the other major ACC output to the dorsal thalamus, where excitation was dominant and inhibition marginal (**Figure S3**).

Next, we dissected how putative excitatory and inhibitory claustrum neurons respond to ACC input by classifying the claustrum neurons described above. We used spike shape to classify single units into regular spiking (RS, putative excitatory neurons) and fast-spiking (FS, putative inhibitory interneurons) as done previously^38,52–54^. RS cells provide the main claustrum output neurons, whereas FS cells mainly project locally within the claustrum^38^. Optogenetic activation of the ACC resulted in the inhibition of most RS cells in the ipsilateral claustrum (61% suppressed, 31% activated) (**Figure 2A-C**) whereas the majority of RS cells in the contralateral claustrum were activated (28% suppressed, 58% activated) (**Figure 2D-I**). To further compare the RS firing response in each hemisphere, we calculated the optogenetic modulation index, the maximum number of spikes per light pulse, the mean firing rate during the 20Hz stimulation, and the entrainment of each cell to the stimulation frequency (**Methods and Materials**). Contralateral RS cells had a significantly greater modulation index, more action potentials per pulse, a higher mean firing rate during the 20Hz ACC stimulation, and greater stimulation entrainment (**Figure 2J-M**). Therefore, ACC inputs drive more excitatory responses in RS cells of the contralateral claustrum relative to the ipsilateral claustrum. Further dissection of RS neurons showed that the degree of excitation in the contralateral hemisphere was greater than in the ipsilateral claustrum (**Figure S4**). Analysis of sex differences in RS responses showed the ipsilateral versus contralateral differences were most prominent in male mice (**Figure S5**). Overall, activity in ipsilateral RS claustrum neurons is predominantly suppressed or weakly activated suggesting ipsilateral claustrum outputs are largely attenuated during ACC stimulation.

**Figure 2:**
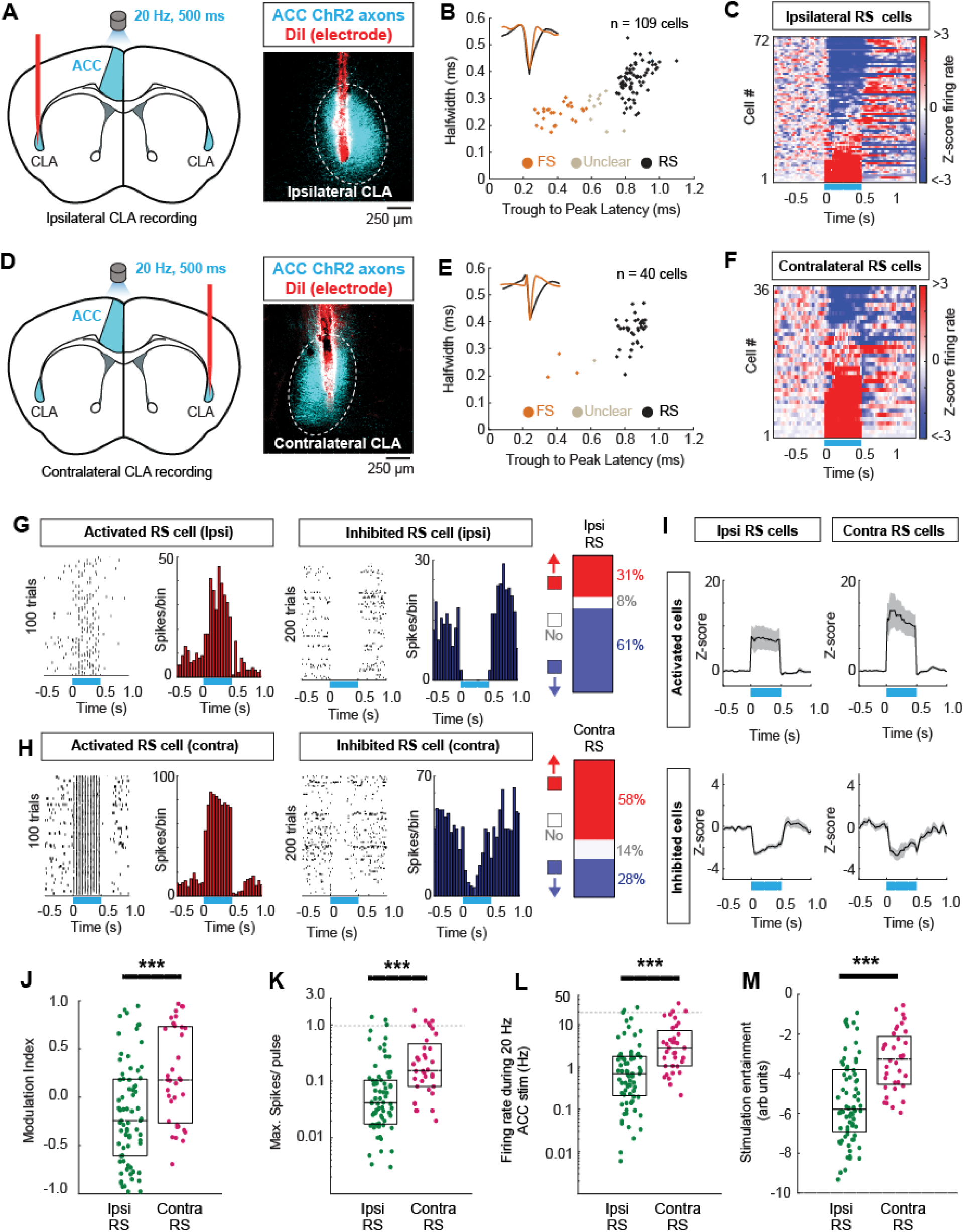
The anterior cingulate cortex drives ipsilateral inhibition and contralateral excitation of claustrum excitatory cells. A: Schematic diagram of ACC optogenetic activation with 20Hz light pulses for 500ms (10 pulses and single unit recordings were made from the ipsilateral claustrum (left). Example electrode and viral expression histology (right). B: Classification of regular spiking (RS, putative excitatory neurons) and fast spiking (FS cells; putative inhibitory neurons). C: Z-scored firing rate for all RS cells recorded in the ipsilateral claustrum in response to ACC activation. D-F: The same as A-C for the contralateral claustrum. G: Example PSTHs showing the firing of ipsilateral claustrum cells that were excited or inhibited as a function of time relative to ACC activation at 20 Hz (blue line). The proportion of activated (red), inhibited (blue), or unmodulated (white) RS cells are shown as well. H: The same as G, but for contralateral claustrum RS cells. I: The average z-scored PSTHs for ipsilateral RS cells (left) and contralateral RS cells (right). J-M: The group data for all ipsilateral and contralateral RS cells. The modulation index (J: Ipsi = -0.24, Contra = 0.18, z = 3.47, p = 5.2×10^-4^), maximum spikes/pulse (K, Ipsi = 0.04, Contra = 0.16, z = 4.56, p = 5.2×10^-6^), firing rate during rate stimulation (L, Ipsi = 0.69Hz, Contra = 2.81Hz, z = 4.93, p = 7.8×10^-7^), and the stimulation entrainment (M, Ipsi = 0.003, Contra = 0.039, z = 4.63, p = 3.6×10^-6^). ***P<0.001.

FS cells in the ipsilateral claustrum were frequently recorded, the majority of which were strongly activated by 20Hz ACC input (**Figure 3A-D**). In total, 20/27 FS cells were activated and 10/27 ipsilateral FS cells showed supralinear response to ACC inputs (firing more than one spike in at least one of the 10 pulses). 6/27 FS cells fired above the 20Hz stimulation frequency showing that a substantial proportion of FS interneurons are responding with high fidelity to ACC inputs. In the contralateral claustrum only 4 FS cells were recorded, and one of these was activated (not shown). For this reason, we did not analyze contralateral FS cells further. To compare RS and FS cells within the ipsilateral claustrum, we first analyzed the latency to activation, specifically in cells that fired within 20ms of the light pulse (and are thus more likely to receive direct ACC input). Activation in FS cells occurred earlier than RS cells (**Figure 3E,F**) in accordance with FS-interneurons having a faster excitatory synaptic response^38,39^. FS cells exhibited a higher modulation index, an increased number of action potentials/light pulse, greater firing rate during the ACC stimulus and increased entrainment, relative to ipsilateral RS neurons (**Figure 3G-J**). Therefore, high fidelity early activation of FS cells is likely preventing the firing of excitatory claustrum neurons through feed-forward inhibition.

**Figure 3:**
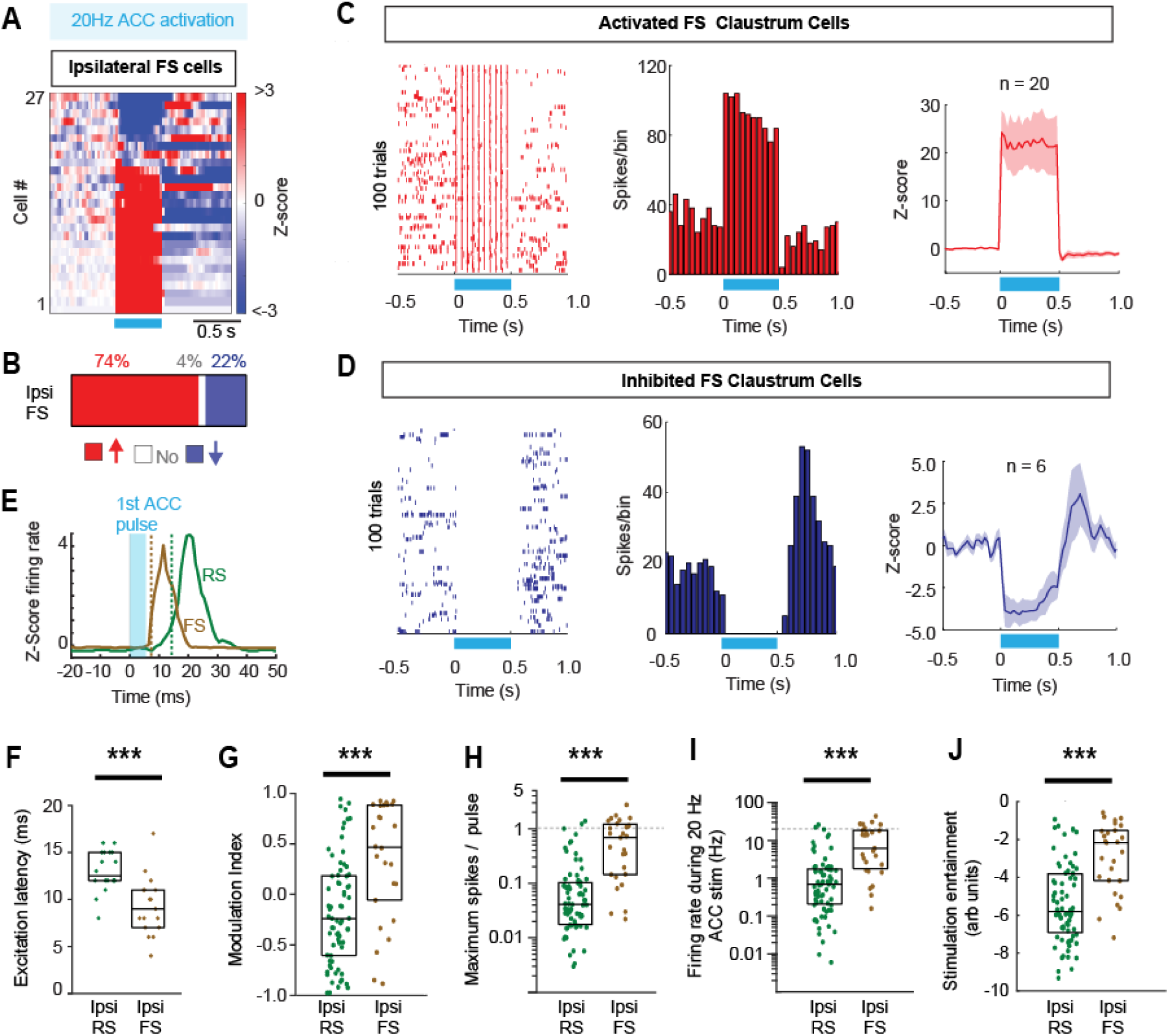
Fast spiking interneurons are rapidly and robustly activated by anterior cingulate cortex activation. A: Z-scored firing rate modulation of all FS interneurons recorded in the ipsilateral claustrum. B: the proportion of FS cells activated, inhibited, or not modulated C: Example spike time raster and PSTH for an activated FS claustrum cell, and the mean z-scored firing across all activated FS cells D: The same as C for inhibited FS cells. E: An example PSTH showing the relatively early activation of an FS cell relative to an example RS cell. The vertical dashed line denotes the activation latency. F-J: The individual FS cells and ipsilateral RS cells (as shown in Figure 2) for each quantified parameter. Latency to excitation (F: RS:13ms, FS:9ms, z = 3.52, p = 4.28×10^-4^), Modulation index (G: RS = -0.24, FS = 0.37, z = 3.69, p = 2.2×10^-4^), maximum spikes per pulse (H: RS=0.04, FS=0.69, z = 5.39, p = 7×10^-8^), firing rate during stimulation (I: RS = 0.69, FS = 6.20, z = 4.94, p = 7.9×10^-7^) and the spike – stimulation entrainment (J: RS = 0.003, FS = 0.11, z = 4.63, p = 3.6×10^-6^). For the latency values, only cells that increased their firing within 20ms of the stimulation were used. Entrainment values were log transformed for display. Box plots show median and interquartile range. Horizontal dashed lines (H and I) indicate the values expected if claustrum neurons responded linearly to stimulation. ***P<0.001.

FS cells are parvalbumin (PV) expressing interneurons, and previous in vitro work has shown that PV cells in the claustrum are activated by ACC inputs^38,39,49^. To determine if claustrum PV interneurons provide ipsilateral feed-forward inhibition, we suppressed claustrum PV interneurons by injecting the inhibitory receptor AAV5-DIO-hM4DGi-mcherry in the claustrum of PV-cre mice. Then, we recorded RS neurons in the claustrum before and after the injection of hM4DGi receptor agonist clozapine-n-oxide (CNO) to reduce the activity of PV interneurons (**Figure 4A-B**). PV neuron suppression increased the ACC evoked activity in the claustrum whereas in control mice no change was detected following CNO (**Figure 4 C-G**). Observation of the stimulation time-course suggests PV cells are contributing more to claustrum inhibition in the early pulses of the ACC stimulation (**Figure 4D-E**).

**Figure 4:**
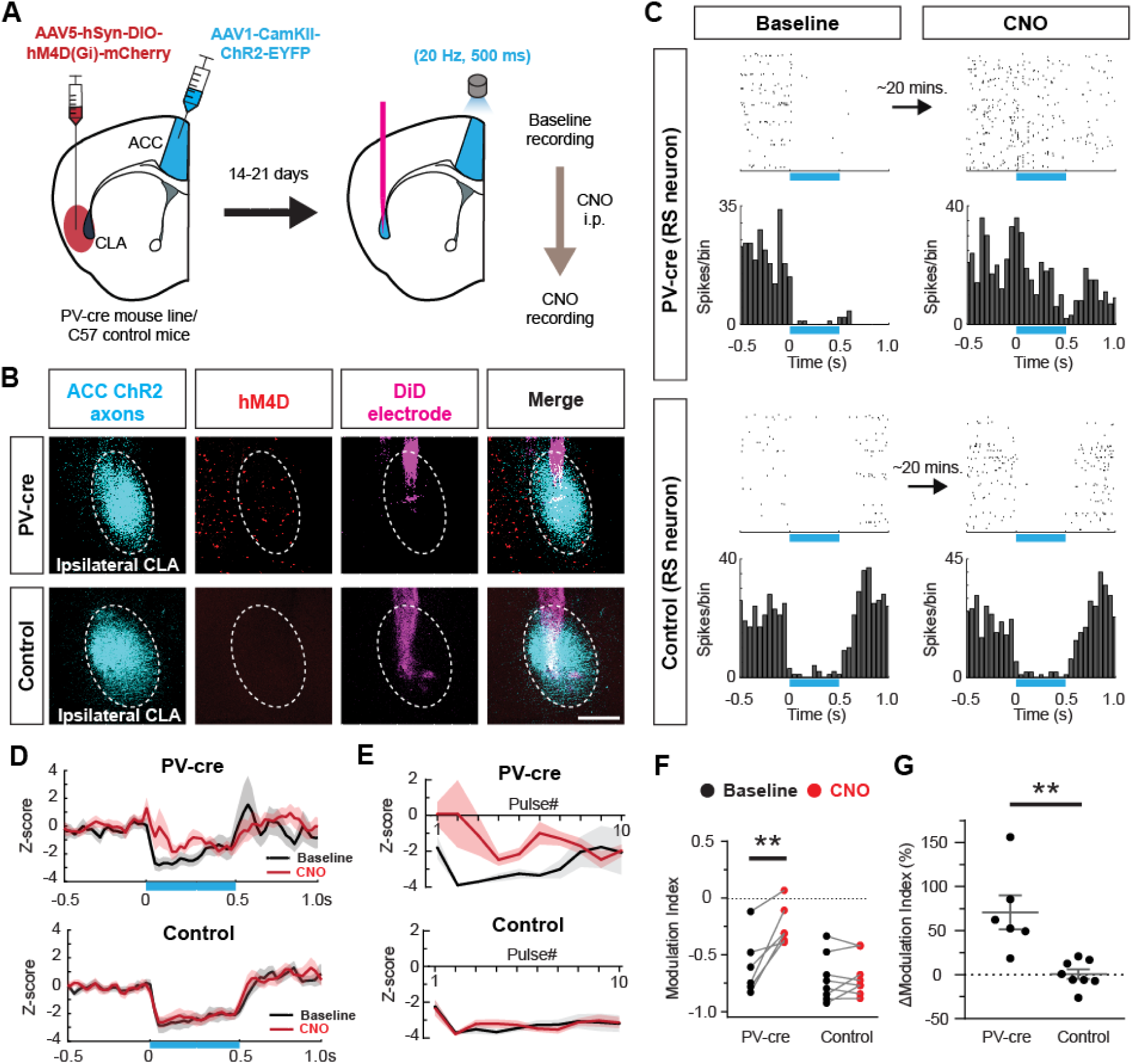
Chemogenetic suppression of claustrum parvalbumin interneurons reduces feed-forward inhibition induced by the anterior cingulate cortex. A: Schematic of the experimental setup showing parvalbumin interneurons in PV-cre mice which were transduced with hm4DGi to suppress PV interneuron activity. Control mice (not expressing cre) were also injected with the same virus. B: Example of post-experiment localization of the electrode in PV-cre mice and control mice (Dil, magenta), showing hm4D(G_i_) expression in PV-cre mice but not in control mice (hM4D, red) relative to the claustrum (ACC ChR2 axons, cyan). C: Example spike rasters and PSTHs where RS cells were recorded in the claustrum prior to (left) and following the administration of clozapine N oxide (CNO, right). Only the cell in the PV-cre mouse exhibits an increase in firing. D: The averaged (±standard error) z-scored firing rate of ipsilateral claustrum neurons before and after CNO administration in PV-cre mice and control mice that do not express hm4DGi. E: The detailed z-scored spiking rates from (D) but measured for each pulse in the 10-pulse sequence of ACC stimulations. Note the impact of PV suppression is predominantly in the first few pulses. F-G: Group data showing the results of all experiments conducted in PV-cre (n = 6 neurons from 6 mice) and control neurons (n = 8 neurons, from 4 mice). The modulation index (F) and the change in modulation index (G: PV-cre = 71±19%, control = 1±5%, p = 0.001) are shown. Each point is one cell. **P<0.01.

To further study the role of PV interneurons in the ACC-claustrum circuit we turned to postnatal development, as it was previously reported that claustrum PV expression emerges after the 2^nd^ postnatal week^55^. The anatomical connections of the claustrum with the ACC appear to be present within the first week of life^51^ but the physiology of the ACC-claustrum connection has not been studied in development. To anatomically label the claustrum at different postnatal ages, we injected retrograde tracers into the ACC or used Nr2f2 and Nurr1 immunohistochemistry^51^ to locate the claustrum and measure the density of interneuron subtypes (**Figure 5A-E, Figure S6**). Indeed, PV cells could not be detected before postnatal day 15 (P15) using either PV-cre::Ai14 reporter mice or immunohistochemistry, whereas somatostatin neurons were present in similar densities across all ages measured (**Figure 5F-G**). Strong PV cell labelling was detected in the overlying somatosensory cortex at P14 (**Figure S6**) indicative of relatively late PV maturation in the claustrum. At P15, some faint PV expression could be detected in PV-cre:Ai14 mice (not using immunohistochemistry), but consistent/reliable (across mice) detection of PV cell bodies and neuropil did not arise until ∼ P17 (**Figure 5H**).

**Figure 5:**
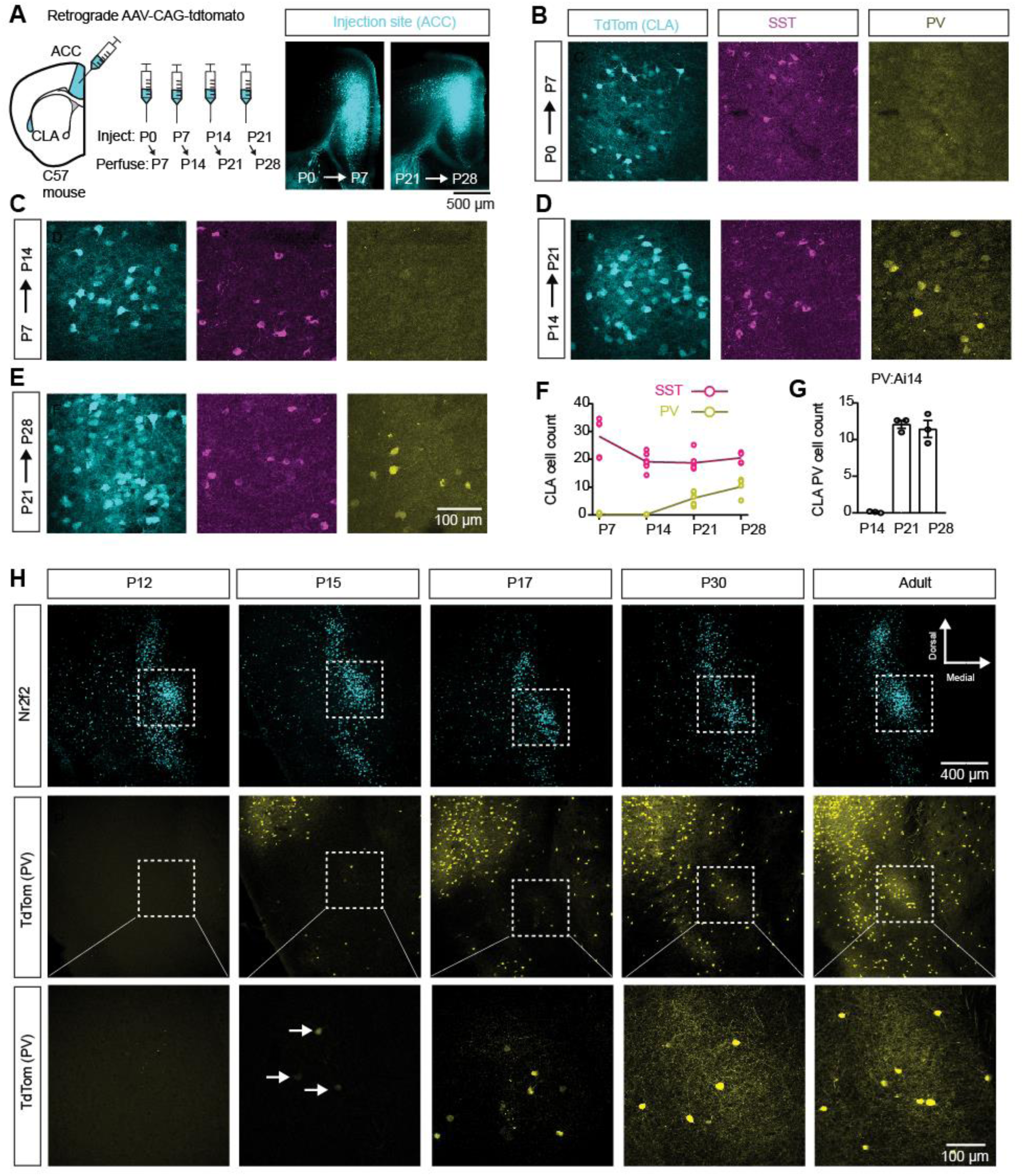
Mature parvalbumin expressing interneurons in the claustrum emerge in the third postnatal week. A: Schematic showing the injection of the retrograde tracer into the ACC. Mice were injected at different postnatal ages and perfused 7 days later. The tracer was used to spatially locate the claustrum, in order to measure parvalbumin (PV) and somatostatin (SST) interneurons at different ages. B-E: Example retrograde labelling of claustrum neurons projecting to the ACC and immunohistochemical labelling of SST and PV interneurons at different injection/perfusion days. F: The group data of PV and SST cell counts on each postnatal day. PV cells could not be detected prior to or at postnatal day 14. P7, n = 5; P14, n = 7; P21, n = 7; P28, n = 4. G: PV neurons were counted in the claustrum of PV-cre::Ai14 reporter to genetically label PV cells. Similar to the immunohistochemical PV labelling, no PV cells could be detected at or before P14 (n = 3 mice/age). H: Example images from PV-cre::Ai14 mice at different postnatal ages. Nr2f2 is used to as a claustrum marker. Nr2f2 is also found in neighboring brain regions, but the densest expression is in the central zone of the claustrum^50,51^. PV cells could be faintly detected on postnatal day 15, and strongly on postnatal day 17 (n = 2-3 mice/age)

To better understand how the ACC drives excitatory and inhibitory signaling in the claustrum prior to PV cell maturation, we performed in vitro patch clamp recordings of claustrum RS cells during optogenetic activation of ACC inputs (**Figure 6A**). Recordings were performed from claustrum RS cells during immature (P14-15) and mature PV timepoints (P42-45) while excitatory and inhibitory synaptic currents were measured in response to ACC input (**Figure 6B**). Optically evoked excitatory postsynaptic currents were larger in young mice while inhibitory currents were equal in magnitude (**Figure 6C**). Excitatory and inhibitory latency and rise times were also not significantly different between the ages. However, the inhibitory decay kinetics were slower in young mice (**Figure 6C**). Faster inhibitory decay is strongly associated with local inhibition arising from FS-PV interneurons^56–59^. Therefore, inhibition is still present before claustrum PV cells mature but is likely mediated by different claustrum interneuron subtypes such as somatostatin neurons which are present in the claustrum prior to PV maturation (**Figure 5F**).

**Figure 6:**
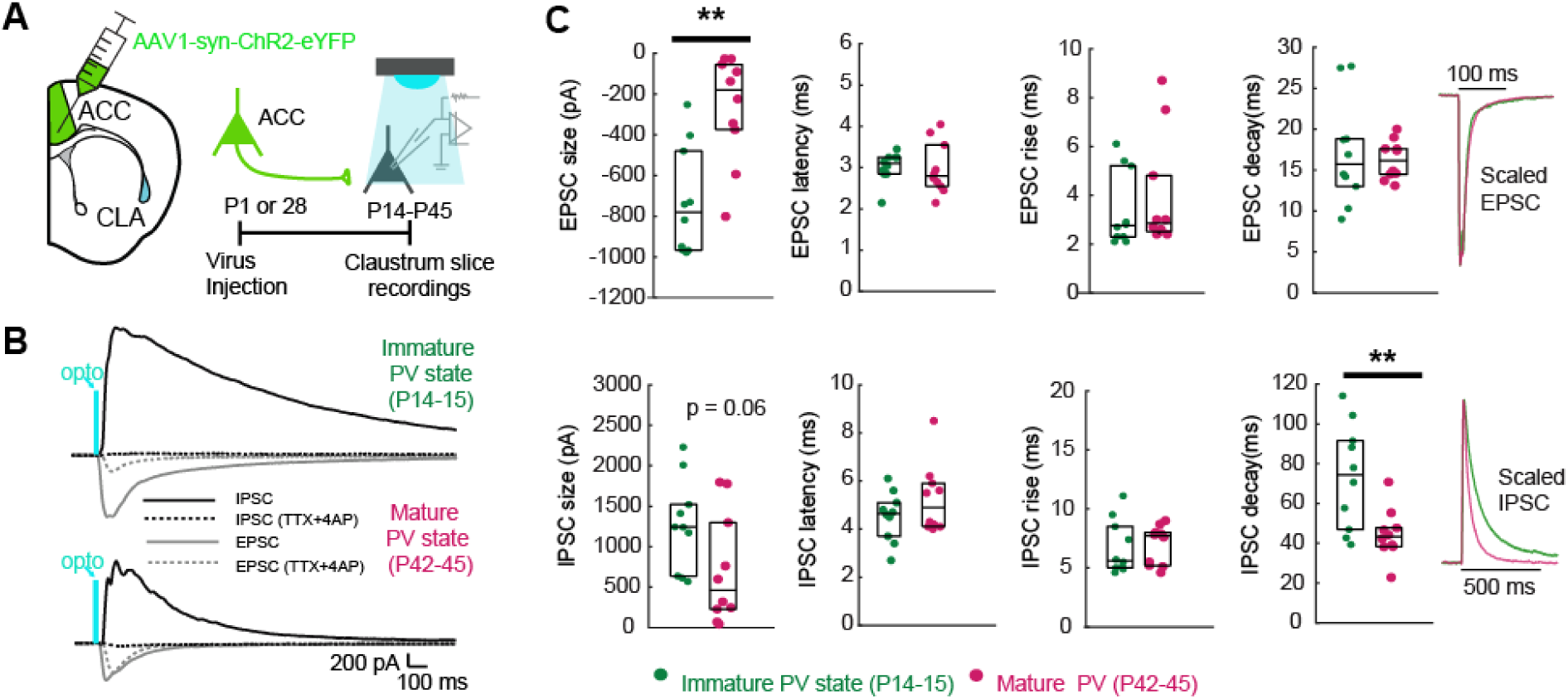
ACC inputs evoke distinct synaptic changes in the claustrum before PV neurons mature. A: Schematic showing viral mediated ChR2 injection in the ACC followed by *in vitro* activation of ACC axons while patch-clamp recording claustrum RS cells in neonatal (immature PV) and adult (mature PV) mice. B: Representative neurons at neonatal and adult time points showing the presence of monosynaptic excitatory post synaptic potentials before and after bath application of 4AP+TTX. C: Group data for EPSC and IPSC size, latency, and decay time at the two times. The mean scaled EPSC and IPSC are shown on the right to highlight the decay. EPSC size: immature PV = - 728±83pA mature PV = -267±83pA, p = 0.001. IPSC decay: immature PV = 73±8ms, mature PV = 45±4ms, p = 0.006).**P<0.01.

These data suggest two possible scenarios regarding the development of ACC-claustrum feed-forward inhibition in the ipsilateral hemisphere. The first is that ACC inputs will act on non-PV cells to generate feed-forward inhibition prior to PV maturation. In this case, ACC activation before PV maturation will evoke suppression of claustrum RS neurons similar to adulthood. The second scenario is that claustrum PV interneurons are critical for feed-forward inhibition and prior to PV maturation ACC activation will evoke excitation in claustrum RS neurons. We performed extracellular in vivo recordings from the claustrum (as described in adults), during optogenetic activation of ACC inputs at P9-22 (**Figure 7A-D**). Optogenetic ACC activation caused excitation in 79% of the RS cells prior to P14 (before PV), and 29% of RS cells at P17+ (after PV) (**Figure 7E-H**). The proportion of inhibitory responses was 13% prior to P14 and 68% from P17 onward. This suggests a transition phase beginning at the third week of life, where the overall response of claustrum RS cells to ipsilateral ACC activation is reminiscent of the adult state. Indeed, the ACC modulation of claustrum spiking shifted to positive values and the number of spikes/optogenetic pulse, stimulation firing rate, and the ACC entrainment were significantly greater before claustrum parvalbumin maturation (**Figure 7I-L**). In addition, whole-cell current clamp recordings in slices showed that P14-15 RS cells fired significantly more action potentials in response to ACC fiber stimulation (0.63 ± 0.51 spikes/pulse, n=16 cells) than did P42-44 RS cells (0.19 ± 0.37 spikes/pulse, n=13 cells, p=0.014, not shown). Therefore, it appears that during maturation, PV interneurons become the dominant source of feed-forward inhibition within the claustrum, as their suppression or absence attenuates the degree of feed-forward inhibition in the claustrum.

**Figure 7:**
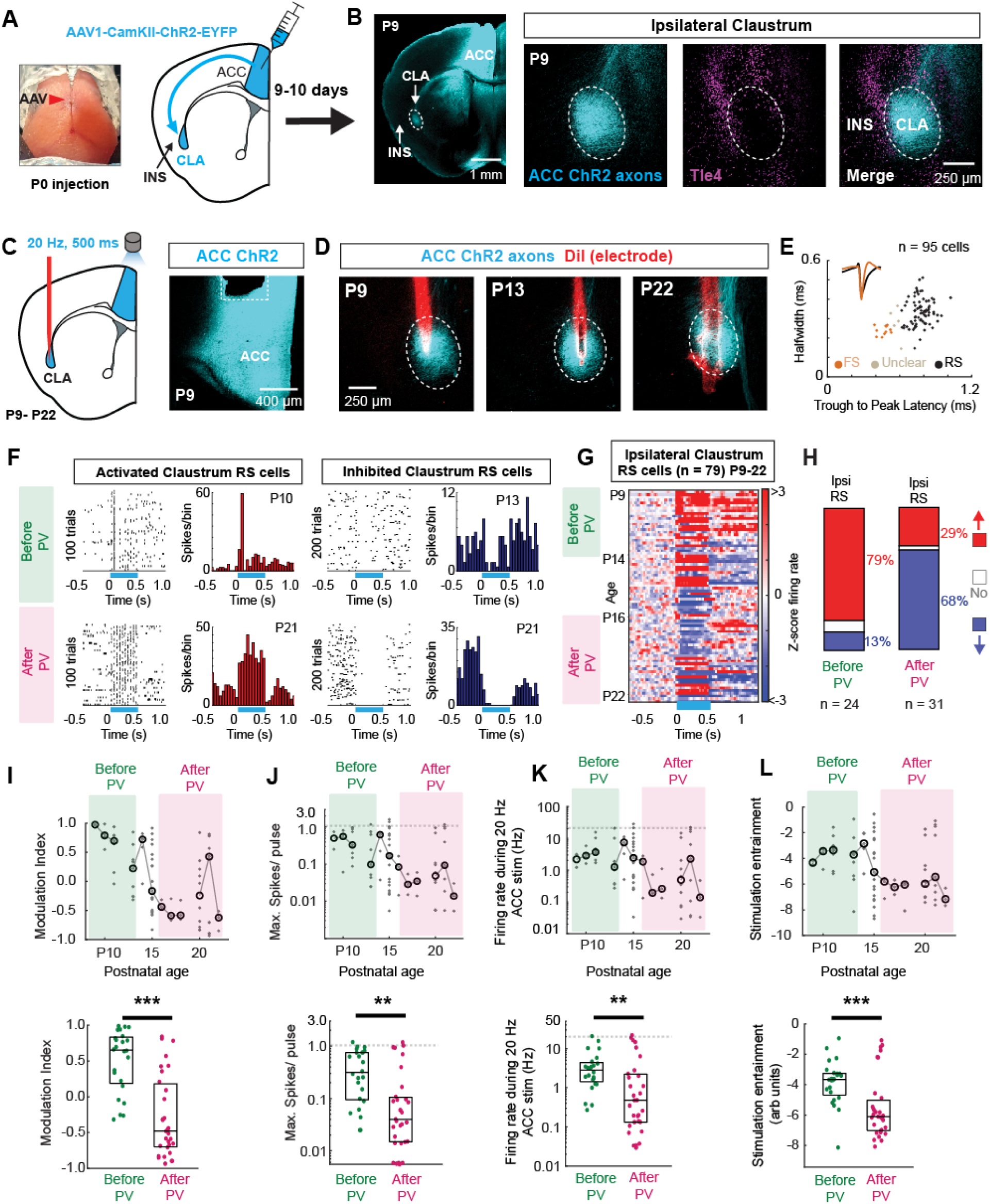
The anterior cingulate activation switches from claustrum activation to claustrum inhibition as a function of PV interneuron maturation. A: Schematic of the injection setup where mice were injected with AAV-CAMKII-ChR2-eYFP at postnatal day 0. B: Histology showing the spread of injection in the ACC at postnatal day 9 (left). Axons and Tle4 labeling in the ipsilateral claustrum at postnatal day 9 (right). C: Schematic showing the recording in the ipsilateral claustrum (left) and histology showing the optic fiber canula (right). D: Examples of post-experiment histology showing the location of recording electrodes in the claustrum at different postnatal ages. E: Cell classification is based on spike waveform characteristics for all cells recorded from postnatal day 9 to 22 (see Methods and Materials). F: Example spike rasters and PSTHs for activated claustrum RS cells (left) and inhibited (right) RS cells during the immature PV state (top) and during PV maturation (bottom). G: Heatmap showing the z-scored firing rate for all 79 RS cells recorded across postnatal development. 25 mice total (mice/group: P9-13: n = 9; P14-15: n=6; P16-22: n=10). H: The proportion of RS cells that were activated, inhibited, or unchanged by ACC stimulation as a function of claustrum PV expression. I-L: The plot of how each claustrum RS cell was changed by ACC activation, as a function of postnatal age. The top shows all cells and bottom shows group data merged across the two ages (before and after PV expression emerged). I: Modulation index (Before PV: 0.65, After PV: -0.48; z = 4.54, p=5.6×10^-6^), J: Maximum spikes/burst (Before PV: 0.31, After PV:0.04; z = 3.27, p=0.001), K: Stimulation firing rate (Before PV: 2.8Hz, After PV: 0.48Hz; z = 2.98, p = 0.003), L: Stimulation entrainment (Before PV: 0.02, After PV: 0.002, z = 3.40, p = 6.7×10^-4^). **p<0.01, ***p<0.001.

## Discussion

Neural activity in the claustrum is involved in mediating aspects of stress regulation, sleep^20–22,24^, attentional engagement^23,49,60^, and pain processing^31–33^. The strongest long-range input to the claustrum arises from frontal midline cortical regions including the ACC^11,34^. Therefore, it is essential to understand how ACC neurons control the activity of claustrum neurons. The data shown here provide the first direct in vivo investigation of how ACC activity controls the activity of different claustrum cell types, complimenting and extending previous in vitro work^30,38–40,42,49,52^.

The key finding here is that in adult mice, the ACC exerts a dissociable change in neural activity in the ipsilateral versus the contralateral claustrum. Inputs from the contralateral ACC show depressing excitatory synapses, whereas ipsilateral inputs are facilitating^39^. Previously, it was shown that the ACC input drives polysynaptic feed-forward inhibition in the contralateral ACC via the activation of the claustrum^30^. This was proposed to help highlight the sensory response to the ipsilateral hemisphere by functioning to suppress activity in the contralateral hemisphere. Our data is in line with this result. The suppression of the ipsilateral claustrum by ACC inputs was more surprising but could further help ensure ipsilateral sensory-motor cortical communication is not interfered with by claustrum inputs. Claustrum inputs to the cortex are most active during offline states such as a slow wave sleep^21,22,61^. Therefore, the ACC-mediated suppression of claustrum-cortical projections within an ‘active’ hemisphere during ‘online’ states of high sensory-motor processing would be advantageous to prevent claustrum signals from interfering with cortico-cortical processing. Other clear instances of claustrum lateralization of activity have been reported in mice and humans. Indeed, it was noted that claustrum activity shows lateralized activity during a multisensory detection task^15^ and pain processing^33^, suggesting that this ACC-claustrum circuit could function across an array of sensory-motor behaviors or even during sleep^62^.

Another key result is the observation that ACC inputs only modestly activate claustrum circuits in vivo and likely fail to relay information arising from the ACC. Previously, it was shown that ACC inputs drive the firing of both excitatory and inhibitory claustrum cells. It was noted that feed-forward inhibition constrained the firing of excitatory cells in *in vitro* experiments, but claustrum neurons were capable of supralinear firing at higher frequencies (>50Hz) far higher than the ACC input frequencies^41,49^. This feature of cortical – claustrum connections helped provide rationale for the theory that claustrum cells integrate and amplify cortical inputs and broadcast this transformed signal back to the cortex. In contrast, our data suggests that claustrum cells in vivo do not faithfully follow the population firing of the ACC. While 5/108 RS cells fired at >20Hz, the median firing rate of RS cells in the contralateral claustrum was only 2.8Hz. The reason for this in vitro/in vivo difference could be explained by the increased preservation of inhibitory connectivity in vivo, or differences in where the optogenetic activation was carried out (axon terminals versus cell bodies). Further study is required to determine if the simultaneous excitation from multiple convergent pathways would be sufficient to enhance claustrum activation, or if the population firing rate would continue to be constrained by feed-forward inhibition.

Our data suggest that the ACC exerts a hemispheric specific modulation of claustrum that correlates with the maturation of claustrum PV expression during postnatal development. However, there is still functional inhibition in the claustrum prior to PV maturation, likely served by other interneuron subtypes which are present (for example somatostatin cells). Indeed, in young mice, cortical inputs evoked inhibition that had slower kinetics, consistent with non-PV interneuron mediated inhibition. These data suggest that as claustral PV cells mature, they become more dominant in mediating ACC evoked feed-forward inhibition. Three separate results support this conclusion. First, these cells fire with an early onset high fidelity response to ACC inputs positioning FS cells to inhibit large populations of excitatory claustrum cells. Second, chemogenetic suppression of claustrum PV cells in adults reduced ACC-mediated feed-forward inhibition. The remaining inhibition after PV suppression likely arises from the imperfect chemogenetic inhibition, and/or from other local claustrum inhibitory cells being recruited by ACC stimulation. Third, PV neurons are first observed in the claustrum at the developmental time point when ACC circuits switch from predominantly evoking excitation to inhibition. Additional interneuron subtypes may also play a role in shaping aspects of ACC mediated inputs. We note that PV suppression in adults (Figure 4) did not recapitulate the degree of excitation found early in development, suggesting that other circuit elements are undergoing changes during early life. One such change is the reduction in excitatory current magnitude seen in vitro (Figure 6C). Collectively, there are marked differences in how these circuits communicate as a function of age, due in part to the development of claustrum inhibition.

PV interneurons migrate into the cortex during the first postnatal week^63^. The expression of PV itself begins to increase in the second postnatal week within the cortex and continues to increase during the third postnatal week^64–67^. Claustral PV expression arises later, as PV immunoreactivity was absent at P14. As PV interneurons enable rapid local circuit oscillations^68–70^ the temporal sequence of cortical PV maturation followed closely by claustral PV maturation suggests that development of oscillatory activity in the claustrum may, to some extent, dependent on oscillatory activity in the cortical circuits that drive it. As PV levels are known to change as in indication of learning and plasticity^71^, this process in the claustrum may be related to sensory-motor learning that arises during the third postnatal week.

Some limitations are inherent in our study. First, the use of extracellular recordings and spike shape for determining cell types must be interpreted with caution. Although narrow spiking neurons are almost always found to be PV positive, the wider spiking RS cells could be either excitatory projection cells or non-PV interneurons. Therefore, we likely overestimate the number of true excitatory cells that are activated by the ACC. The second limitation is that the projection target of the RS cells we recorded is not precisely known. Our dataset was limited to those recordings that were histologically confirmed to be within the anterior claustrum, as defined by ACC axon innervation pattern like previous reports. This region of the claustrum has excitatory cells projecting to the ACC, the retrosplenial cortex, orbitofrontal cortex, and secondary visual cortex. Future experiments using optogenetics to tag specific claustrum outputs^15^ would be useful to determine the projection target of the recorded neurons.

To conclude, we provide vivo data showing how the ACC communicates with the claustrum, demonstrating a PV-dependent feed-forward inhibition that develops in the third postnatal week. The hemispheric specific modulation could provide an efficient long-range circuit by which the ACC can down-regulate the activity in the contralateral hemisphere using claustrum mediated feed-forward inhibition^30,48,54,72–74^. At the same time, the suppression of the ipsilateral claustrum would ensure claustrum activity does not interfere with ipsilateral top-down signals from the ACC to sensory cortex. ACC activation has been shown to enhance attention^3,49,75^. Therefore, it is possible that suppression/activation of the ipsilateral/contralateral claustrum participates in balancing hemisphere specific excitability of cortical circuits required for cognitive demands.

## Materials and Methods

### Experimental animals

Animal care and experimental procedures were approved by the University of Alberta Animal Care and Use Committee (AUP-3715) and were in compliance with the Canadian Council on Animal Care Guidelines. Wild type and transgenic mice were maintained on a C57BL/6 background. Ai14 (Jackson Laboratories Stock #007909^76^), and, PV-cre (Jackson Laboratories stock # 017320) transgenic mouse lines were used. Both sexes were included in the study. Mice between 55 and 120 days old were used for adult experiments, and mice between 0, i.e. newborn, and 22 days old were used for developmental experiments. Mice were housed in temperature-controlled conditions with a 12-h light/dark cycle, and with ad libitum access to food and water.

### Stereotaxic injection of viral vectors

Adeno-associated viruses (AAVs) were purchased from Addgene (MA, USA) at titer ≥ 1×10^12^ vg/ml. pAAV-CaMKIIa-hChR2(H134R)-EYFP was a gift from Karl Deisseroth (Addgene plasmid # 26969 ; http://n2t.net/addgene:26969 ; RRID:Addgene_26969)^77^. pAAV-CAG-GFP was a gift from Edward Boyden (Addgene viral prep # 37825-AAVrg.T; RRID:Addgene_37825). pAAV-CAG-tdTomato (codon diversified) was a gift from Edward Boyden (viral prep # 59462-AAVrg; RRID:Addgene_59462). pAAV-hSyn-DIO-hM4D(Gi)-mCherry was a gift from Bryan Roth (Addgene viral prep # 44362-AAV5); RRID:Addgene_44362)^78^. pAAV-hSyn-Cre-P2A-dTomato was a gift from Rylan Larsen (Addgene viral prep # 107738-AAVrg; RRID:Addgene_107738). Injections at P0-P21 and adult stages were unilaterally performed as previously reported^51^. Briefly, at P0, pups were cryo-anesthetized and placed in a stereotaxic apparatus equipped with a homemade head holder. A glass pipette connected to a micropump (Narishige, Japan) was lowered into the ACC (A/P: + 2.35, M/L: ± 0.20, D/V: − 0.65), and 75-100 nl of undiluted AAV was injected at a rate of 50 nl/min. A/P coordinates at were relative to the intersection of the transverse sinus and the superior sagittal sinus. Following injection, the pipette was left in place for 1 min, then brought up to D/V: − 0.35 and left there for 1 min, before the pipette was completely withdrawn. Pups were placed on a heating blanket, and once fully recovered, they were returned to their homecage. We followed the same experimental procedures for P7-21 and adult mice, except that P7-21 subcutaneously received carprofen (2 mg/kg) 5–10 min before starting the surgery, whereas adult mice were given carprofen (5 mg/kg), from 24 h before surgery. For adult mice, they were given carprofen (5 mg/kg), from 24 h before surgery, and until 72 h after surgery.

Mice were operated under isoflurane anesthesia (induction 3.5-4%, maintenance 1.5-2.0%). The head was fixed in a stereotaxic apparatus while resting on a heating pad. After exposing the scalp, Bupivacaine was applied, and an incision was made under sterile conditions. A craniotomy was performed with a motorized drill, and a glass pipette was lowered to the target site. The following coordinates were used: ACC (P7, A/P: + 0.40, M/L: + 0.25, D/V: − 0.30; P14, A/P: + 0.65, M/L: + 0.30, D/V: − 0.40; P21 and adult mice, A/P: + 1.00, M/L: ± 0.45, D/V: −1.00), CLA (adult mice, A/P: + 1.50, M/L: ± 2.75, D/V: − 2.60). A/P coordinates were relative to bregma and D/V coordinates were relative to brain surface. For optogenetics, 200 nl of undiluted AAV was injected in the ACC for adult mice (75-100nl at P0), whereas for chemogenetics, 350 nl of AAV diluted with equal volume of sterile PBS was injected in the claustrum/insula region. The large AAV volume in chemogenetic experiments transduced cells in other brain regions, but the manipulation and recording should be specific for claustrum circuits given that the ACC axons do not substantially innervate brain regions surrounding the claustrum.

### In vivo electrophysiological recordings

Mice injected with AAVs were used for electrophysiological recordings in the claustrum, insula, or thalamus. Recordings in neonatal pups were made between P9 and P22, and in adult mice 2-3 weeks post injection. On the day of recording, animals were anesthetized, subcutaneously injected with carprofen (2-5 mg/kg), placed in a stereotaxic frame, and the skull was exposed and leveled following the surgical procedures described above. Three craniotomies were made: 1) over the ACC region at the site of AAV injection, 2) above the claustrum/insula region ipsilateral or contralateral to injection site, 3) on top of the cerebellum, and in some mice 4) above the lateral dorsal thalamus ipsilateral to injection site. Coordinates for craniotomies in the ACC and the claustrum/insula were the same as above, and in the thalamus A/P: -1.6, M/L: ± 1.0. A reference electrode (with an impedance similar to the recording electrode) was placed above the cerebellum, between skull and the dura, and secured with Vetbond tissue adhesive (3 M, MN, USA) followed by dental cement. After thoroughly drying the skull surface, a head plate was attached to the skull, and the skull was sealed with quick adhesive cement (C&B Metabond, Parkell Inc., NY, USA), with the craniotomies left accessible for recording. While the mouse was still under anesthesia, it was quickly transferred to a recording chamber equipped with an isoflurane vaporizer and a heating pad, and the headplate was firmly fixed to a headstage. Using a stereotaxic micromanipulator (Narishige, Japan), a cannula (Ø1.25 mm ferrule, 200 µm core, 0.39NA, RWD) attached to a single-mode optic fiber (200 µm core, 0.39NA, Thorlabs, NJ, USA) was inserted into the craniotomy designated for the ACC, and a 4 channel silicon microelectrode (Q1×4-10mm-50-177-Q4, NeuroNexus, MI, USA) was lowered into the claustrum/insula region, and in some cases into the thalamus, at a low speed to minimize tissue damage. Each channel on the probe were 0.1mm apart. Craniotomies were kept covered with 0.9% saline. The cannula and the electrode were left in position for at least 20 mins before starting the recording session. Knowledge of the orientation and channel geometry of the multichannel probe enabled the determination of the location of the recorded neuron to ∼ 20-50µm. For all our recordings the active channels on the probe were orientated to face midline.

For optogenetic stimulation, light was emitted from a 473 nm laser (DPSS Laser System, CA, USA at ∼7mW intensity measured at the tip of the optic fiber. Trains of 5ms light pulses were delivered using Master-8 (AMPI, Israel) at a frequency of 20 Hz (10, 5ms pulses for 500 ms). 20Hz stimulation was chosen for several reasons. First, previous work has used this frequency to study ACC-claustrum physiology in vitro, so this enabled a direct comparison. Second, this brief 20Hz, 0.5s, stimulation enabled enough time to study short-term plasticity (facilitation or depression) of spiking if it were present. Third, this brief burst of ACC activity is likely modelling periods of high ACC engagement in ACC-dependent behaviors. We could not test multiple frequencies, because so many trials are required to properly construct the PSTHs, therefore, a single stimulation frequency had to be chosen. In our experience, the detection of inhibition often required 200-500 stimulation trials (requiring over 1.5 hours/recording) due to the low baseline firing (1-3Hz) of neural activity. Optogenetic activation of ACC projections was verified by detecting local field potential activity in the claustrum/insula region or thalamus. Data was recorded with an OpenEphys acquisition system, filtered between 1Hz-10kHz and acquired at a sampling rate of 30 kHz. Once a recording was obtained, the electrode was withdrawn and re-inserted 50-100 μm posterior to the location of previous recording. For each location, at least 100 trials with ACC stimulation were recorded. Throughout the recording, the mouse was maintained in a lightly anesthetized state by setting isoflurane ∼1% and oxygen at ∼1 L/min. Occasionally, when the mouse wakes up from anesthesia during recording, light pulses are stopped, isoflurane is turned up to 1.25-1.50% for a few minutes, then turned down to ∼1%. Once the mouse returns to the light anesthesia state, light pulses are turned back on, and the recording is resumed. The total amount of time taken to carefully lower the electrode, find cells, and perform enough stimulation trials (100-500) to get enough spikes to properly construct the peristimulus time histograms was ∼ 2 hours. Therefore, to perform multiple recordings in the same mice, we used lightly anesthetized mice to increase the amount of data collected/mouse and to ensure histology on every recording. In addition, recordings in young ∼ P9-22-day old mice required anesthesia because we could not effectively habituate these mice for awake recordings. Our unpublished data show that claustrum activity rates are conserved between awake rest and light ∼ 1% isoflurane anesthetic, making this preparation convenient for studying claustrum physiology.

For chemogenetic experiments, after 3-4 weeks of injecting AAVs mediating hM4D(Gi) expression as described above, ACC optogenetic stimulation and claustrum recording were conducted like all other *in vivo* experiments. Once a claustrum neuron was recorded where ACC stimulation resulted in activity inhibition, clozapine-n-oxide (CNO) (Enzo Life Sciences, NY, USA) suspended in 0.9% saline was administered (2.5 mg/Kg i.p.). After 20 mins, the same neuron was recorded again. All recordings were performed within 1.5 hours of CNO delivery.

Post-mortem histological labeling of the electrode tracks was performed at the end of each session and once all recordings were obtained. The electrode was withdrawn, coated with Dil, or DiD (ThermoFisher Scientific, MA, USA), and re-inserted into the same location as the last recording. After 10-15 mins, the electrode was retracted, the mouse was perfused, and brain tissue was subsequently processed, sliced and imaged as described below. AAV expression at the injection site and target regions were examined in brain slices using immunolabeling as described below. Only mice where the injection was confined to the ACC and the electrode position was confirmed to be in target brain structures, i.e. claustrum, insula and thalamus, were used for further analysis.

### Tissue processing and immunohistochemistry

Mice were transcardially perfused, and brains were extracted, post-fixed in 4% paraformaldehyde and coronally sliced at 100 μm with a vibratome (5100mz; Campden Instruments, UK) as previously described^51^. For fluorescent immunolabeling, slices were first incubated in phosphate buffered saline (PBS) in Triton-X (PBST): 0.3% Triton X-100 in 1 × PBS (pH 7.4), for 10 mins at room temperature (RT), then with blocking solution: 3% Bovine serum albumin in PBST, for 1 hr at RT, followed by primary antibodies overnight at 4 °C. The following primary antibodies were used at the indicated dilutions in blocking solution: Chicken anti-GFP (gfp1010; Aves Labs; 1:1,000), Rabbit anti-Nr2f2 (ab211776, Abcam; 1:250), Goat anti-Nurr1 (AF2156, R&D Systems; 1:250), Goat anti-PV (PVG213, Swant; 1:2,500), Rat anti-SST (MAB354, Millipore; 1:250), Mouse anti-Tle4 (sc365406, Santa Cruz Biotechnology; 1:250). The next day, slices were washed 3x in PBST for 10 mins at RT, then incubated in secondary antibodies for 3 hrs at RT and washed 3x in PBST. The following secondary antibodies were used at 1:1,000 dilution in blocking solution: (ThermoFisher Scientific, MA, USA): Donkey anti-Chicken IgY Alexa Fluor 488 (A-78948), Donkey anti-Chicken IgY Alexa Fluor 647 (A-78952), Donkey anti-Goat IgG Alexa Fluor 647 (A-21447), Donkey anti-Goat IgG Alexa Fluor 555 (A-21432), Donkey anti-Mouse IgG Alexa Fluor 488 (A-21202), Chicken anti-Rat IgG Alexa Fluor 647 (A-21472). Finally, slices were mounted on slides using Prolong Gold (ThermoFisher Scientific, MA, USA).

### Imaging

TCS-SP5 and SP8-STED confocal microscopes (Leica, Germany) were used to acquire all images using a 10x objective (0.3 NA for TSC-SP5 and 0.40 NA for SP8-STED), and 25x objective (0.5 NA TSC-SP5 and 0.95 NA for SP8-STED). Sequential multi-channel acquisition was set for 488, 543 and 633 nm emission, and AF488, AF555 and AF647 default filtering, using 1 airy unit, 400 Hz unidirectional speed, 1,024 × 1,024 pixel format, 1.00 zoom, 2 μm z-step size and 20 µm stack volume. All imaging parameters were identical across all mice. Claustrum images used for quantification were taken from anterior, middle and posterior subdivision with 2 slices/subdivision separated by 100–200 µm in ≥P28 mice and by 50–100 µm in P7-P22 pups.

### In vivo analysis

*In vivo* electrophysiological recordings were analyzed offline using custom scripts generated with MATLAB (Mathworks, MA, USA). For isolating spiking activity, raw data were bandpass filtered at 0.2 kHz - 15 kHz. Units were sorted based on principal component analysis and were deemed to be a single unit only when there was < 0.1% of events within a 2-ms inter-spike interval. Single units with spike threshold < 75uV, and < 1000 events, were excluded from further analysis. Units were sorted using the mixture of skew-t distributions algorithm as described previously^79^. After initial clustering, all units within a recording were correlated against one another based on their mean waveform. Clusters with a correlation coefficient exceeding 0.90 were merged, provided that the merged cluster did not show spikes in the 2-ms inter-spike interval exceeding 0.1% (if the merging did cause this, the unit with more total spikes was chosen and the other was discarded). A time window of 2 seconds prior to the first light pulse was considered the pre-stimulation period, i.e. baseline. Z-score of firing rate (FR) change during the stimulation period (stim) was calculated relative to the mean FR during baseline. Mean z-score during the entire stimulation period (500 ms) was calculated to sort units based on their modulation. The cutoff for change in modulation was manually set at z-score = 0.2. Single-units were considered activated when z-score > +0.2, inhibited when z-score < -0.2 and no response when z-score between -0.2 and 0.2. Modulation index (MI) was calculated as: (FR_stim_ – FR_baseline_) / (FR_stim_ + FR_baseline_). To separate RS and FS cells, units from each experimental group were clustered based on spike half-width and trough-to-peak duration. The cutoff threshold for FS cells was manually set at spike half-width < 0.30 ms and trough-to-peak duration < 0.60 ms. Cells with spike half-width and trough-to-peak duration values close to the FS cell cutoff threshold were clustered separately and were categorized as “unclear”. In all figures, the first 100 trials are shown for activated cells and the first 200 trials for inhibited cells, 50 ms bins were used in PSTH, 25 ms bins for heatmaps and z-score plots, and 1 ms bins for determining spike latency. The maximum spikes/pulse was calculated by first calculating the average number of spikes per optogenetic light pulse for each of the 10 pulses. The maximum value of this set was used. Therefore, this is likely an upper limit for the estimate of spikes per pulse. The stimulation entrainment was calculated by performing a Fourier transform of the 0.5s of the PSTH where the stimulation was delivered and extracting the power at 20Hz. Therefore, the stimulation entrainment value is expressed in arbitrary units that reflect how well spikes were driven at 20Hz.

For histology, confocal stacked images were exported to FIJI (ImageJ, NIH, MD, USA) and flattened using maximum intensity projection. Demarcation of target area together with cell registration for co-localization and quantification analysis were performed using custom MATLAB scripts. Spatial analysis of fluorescence intensity was quantified and plotted in MATLAB (Mathworks, MA, USA) as previously described^44,51^.

### In vitro slice preparation

Mice were injected as described above with 175nl of a 1:10 mixture of AAV1-syn-ChR2-EYFP and AAVrg-hSyn-Cre-P2A-dTomato bilaterally at P1 or P28. Mice were euthanized at P14-15 or P42-44 by cervical dislocation under isoflurane anesthesia. P14-15 brains were quickly removed following decapitation and placed in ice-cold artificial cerebrospinal fluid (ACSF), which included (in mM): 126 NaCl, 3 KCl, 2 MgSO4, 1.1 NaH2PO4, 2 CaCl2, 10 Glucose, 0.1 Na-pyruvate, 0.4 ascorbic acid, 0.5 glutamine, and 26 NaHCO3 (pH 7.3-7.4 and 300-315 mOsm), while P42-44 brains were placed in ice-cold N-methyl-D-glucamine (NMDG) solution, which included (in mM): 2.5 KCl, 1.2 NaH2PO4, 30 NaHCO3, 20 HEPES, 25 Glucose, 5 Na ascorbate, 3 Na pyruvate, 93 NMDG, 10 MgSO4, and 0.5 CaCl2 (pH 7.3-7.4, 300-310 mOsm). Coronal slices (250 μm thick) containing the claustrum were prepared using a vibratome (Leica VS1200). The slices were collected and placed in a holding chamber with ACSF (P14-15) or NMDG solution (P42-44) maintained at 34°C. After 20 minutes, the slices were transferred to another holding chamber containing ACSF and allowed to recover for at least 1 hour at room temperature before recordings. During slicing, recovery, and recordings, the solutions were continuously bubbled with 95% O2 and 5% CO2. To verify viral targeting, slices containing the injection location were post-fixed in 4% PFA for 3-5 days, then mounted with Mowiol mounting medium with 5 μM DAPI for nuclear counterstain. Mice were only included in the analysis if EYFP expression was restricted to the injection location.

### In vitro recording and analysis

For electrophysiological recordings, brain slices were placed in a recording chamber that was continuously perfused with oxygenated ACSF maintained at 32°C. Neurons in the claustrum were visualized using an Olympus BX51WI upright microscope equipped with infrared differential interference contrast optics. Patch pipettes, with a resistance of 3-6 MΩ, were pulled from borosilicate glass capillaries and filled with an internal solution based on potassium-gluconate, containing in (in mM): 130 K-gluconate, 5 NaCl, 2 MgSO4, 10 HEPES, 0.1 EGTA, 4 MgATP, 0.4 Na-GTP, and 2 Pyruvic Acid (pH adjusted to 7.3 with KOH). Recordings were performed using Multiclamp 700B amplifiers (Axon Instruments, Molecular Devices). The data were filtered at 3 kHz, digitized at 10 kHz (using a National Instruments USB-6343 digitizer), and collected with MIES software (https://github.com/AllenInstitute/MIES) running in IgorPro 8 (WaveMetrics). Recordings were included if access resistance (monitored with a 10 mV hyperpolarizing step between stimulation sweeps) was <25 MΩ and did not increase more than 20%, and if the net leak current did not exceed -200pA. To capture locations of recorded neurons an image of pipette placement within the claustrum was captured after each recording with a 4x objective. Response kinetics were measured on averaged traces with baseline from 30 ms pre-stimulus; onset latency was time to 4× baseline SD, rise time from 10–90% peak, and decay time from exponential fit to 36.8% peak.

### In vitro optogenetics

Photostimulation of ChR2 was achieved by delivering 470 nm light through a 40x objective using an LED system (DC4100; Thorlabs Inc, USA). Blue irradiance was set at 0.7-1 mW/mm2. Cells were voltage-clamped at -70 mV to record EPSCs and at 40 mV to record IPSCs, both in response to single 2 ms pulses of 470 nm light. TTX+4-AP were added to test monosynaptic connectivity. Firing rates were assessed in current clamp mode at the resting membrane potential of the cells.

### Statistics

Statistical analyses were performed using GraphPad Prism 8.0 (Dotmatics, MA, USA) or Matlab v2018b. Differences between 2 experimental groups were assessed using unpaired Mann Whitney U test. Data are represented as mean ± standard error of the mean (SEM) or median ± interquartile range, with n being number of individual mice or single unit activity as reported in the figure legends. Differences were considered statistically significant at p < 0.05.

## Acknowledgements

We thank The Faculty of Medicine & Dentistry Cell Imaging Center for support, and Dr. Anna Taylor for comments on the manuscript. Parts of this project have been made possible by the Canada Brain Research Fund (CBRF), an arrangement between the Government of Canada (through Health Canada) and Brain Canada Foundation, and by the Azrieli Foundation. Funding was provided by Grant/Award Number: 37931; Canadian Institutes of Health Research, Grant/Award Number: 426485; Natural Sciences and Engineering Research Council of Canada, Grant/Award Number: RGPIN2018-05212, the Calgary Foundation – Peter Lougheed Research award. JJ holds a Canada Research Chair award.

## Author contributions

Design/Conceptualization: TS, MMK, JJ. Experiments and Methodology: TS, LR. Analysis: All authors. Supervision: MMK and JJ; Writing: TS and JJ. Funding: MMK and JJ.

## Declaration of interests

The authors declare no competing interests.

---

**Figure S1:**
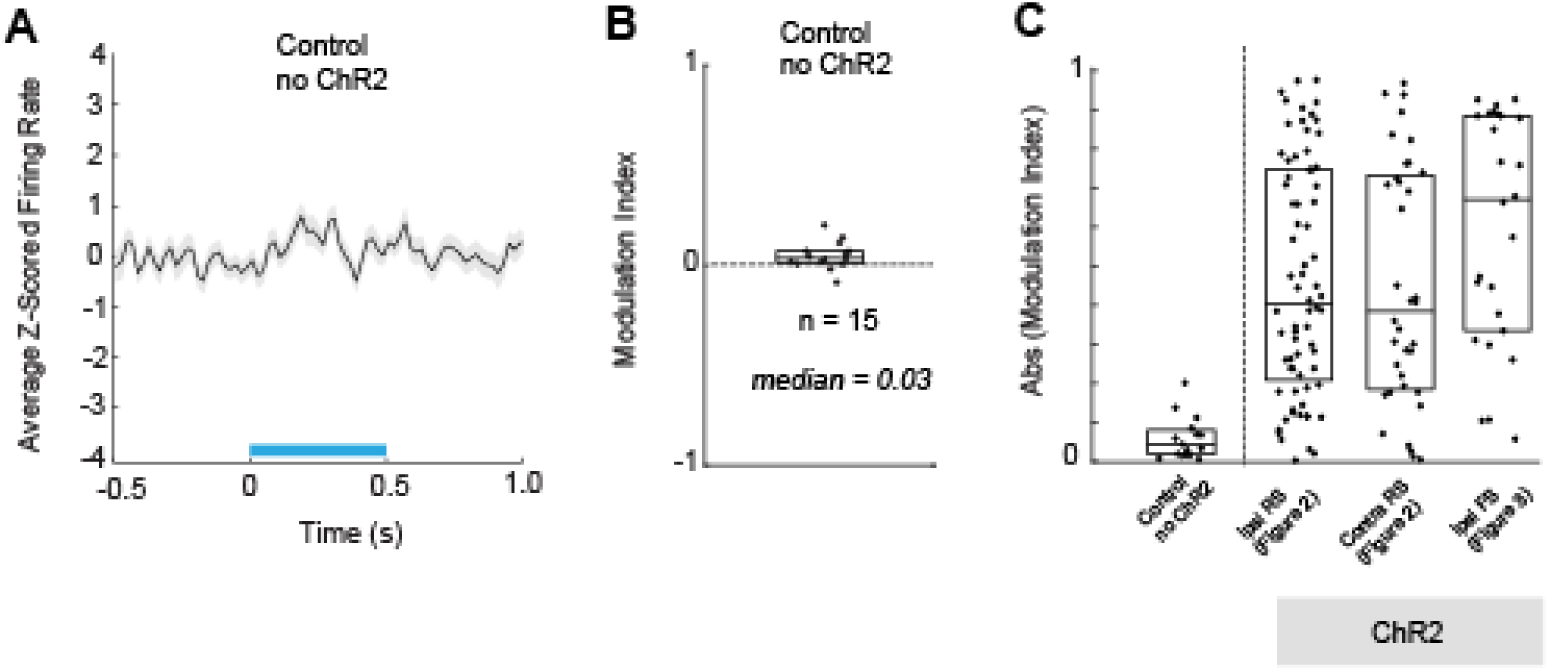
Light delivery to the ACC in control mice does not meaningfully impact claustrum activity. A: The mean Z-scored firing rate for claustrum neurons recorded in control mice that had no optogenetic virus in the ACC. In these mice, light was delivered as in the optogenetic experiments using 20Hz stimulation for 0.5s (blue line indicates stimulation period). B: The modulation index of the cells in these control experiments. C: Plot showing the response of ipsilateral claustrum neurons to ACC stimulation in control mice compared to mice with optogenetic viral injection. Absolute values of the modulation index for cells shown in the main figures are shown here relative to the control. Box plots show the median ±interquartile range.

**Figure S2:**
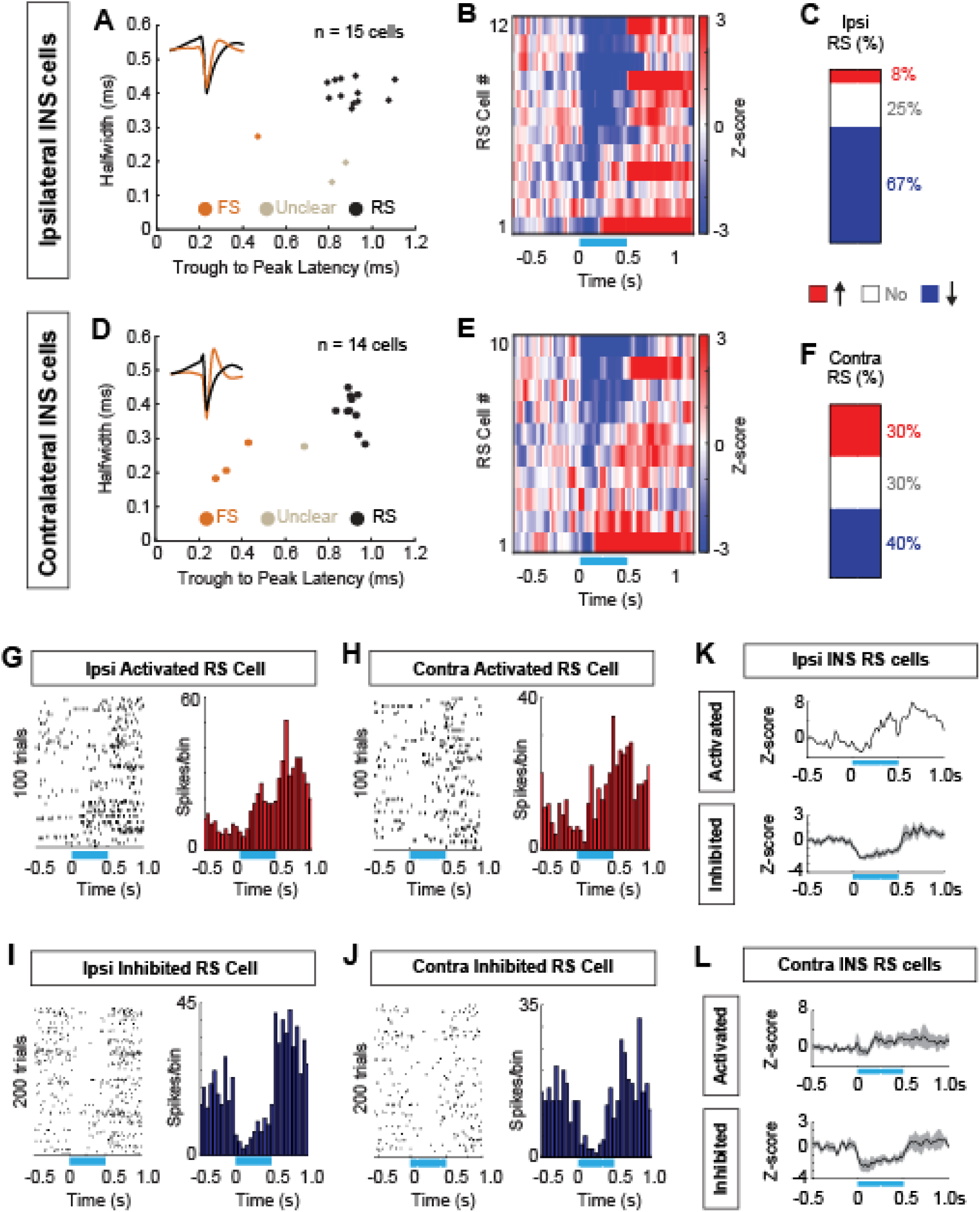
Insula neurons in both hemispheres are mainly inhibited by ACC stimulation. A-C: Data for the ipsilateral insula is shown. Spike waveform properties (A), z-scored firing rates for the RS cells (B), and the percentage of cells that were increased, decreased or unchanged by stimulation (C). D-F: Same as A-C for contralateral insula RS cells. G-H: Example spike raster and PSTH for insula RS cells that were activated by ACC stimulation in ipsilateral (G) and contralateral (H) insula. I-J: Example spike raster and PSTH for insula RS cells that were inhibited by ACC stimulation in ipsilateral (I) and contralateral (J) insula. K: The mean Z-scored firing rate for activated (top) and inhibited (bottom) RS cells in the ipsilateral insula. L: The mean Z-scored firing rate for activated and inhibited RS cells in the contralateral insula. Shaded areas shows the standard error of the mean. Note that in (K, top), variance is not shown because only one neuron was activated in this group.

**Figure S3:**
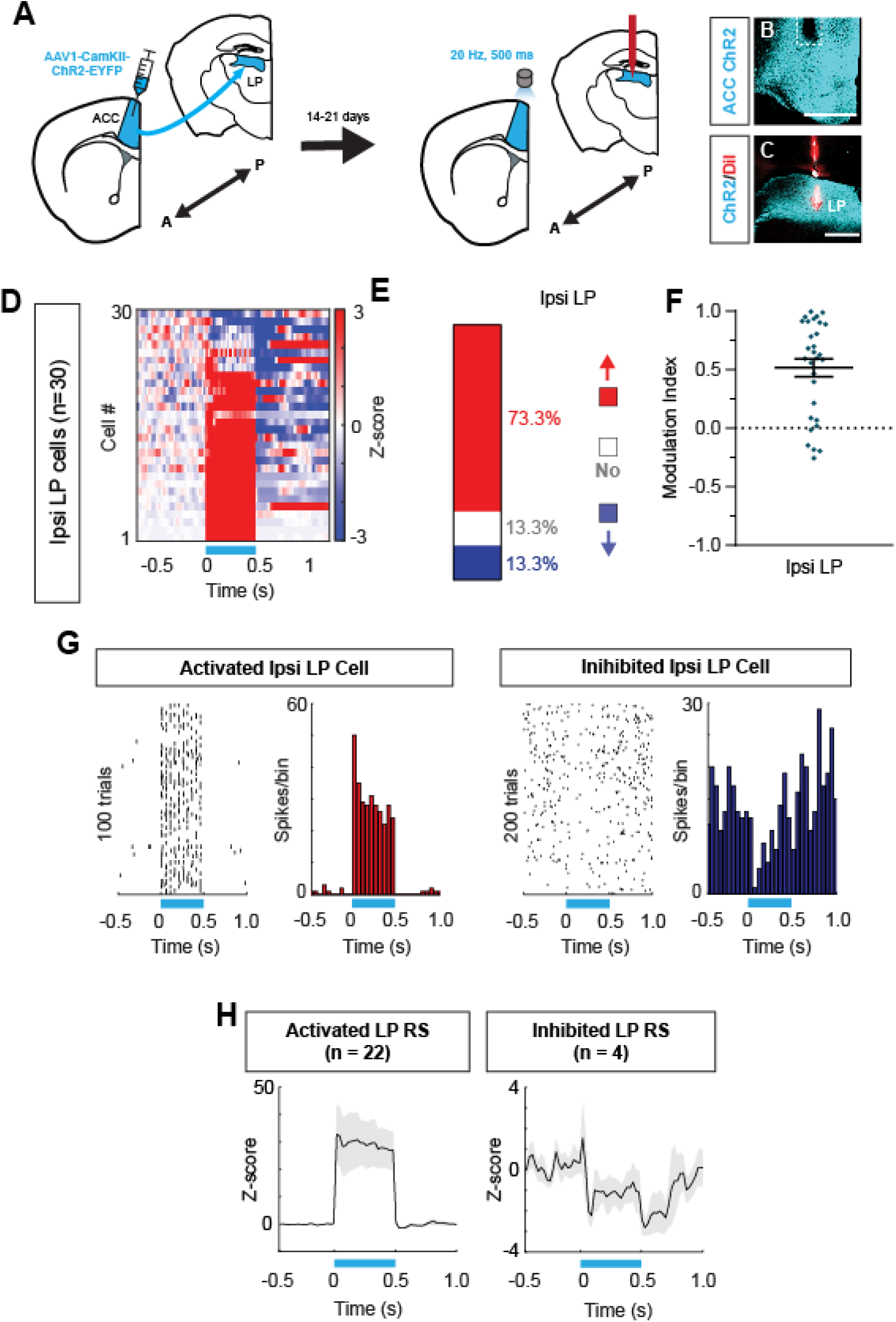
Optogenetic activation of the ACC drives excitation in the thalamus. A-C: Schematic of ACC virus injection followed by *in vivo* recording of cells in the lateral posterior thalamus (LP) during ACC stimulation. Example histology showing post hoc location of the optic fiber in the ACC (B) and recording electrode in the LP (C). Scale bar is 0.5mm. D: The z-scored firing rate for all thalamic cells recorded. E: The percentage of thalamic cells recorded that were activated, inhibited, or unchanged. F: Modulation index for ACC-evoked thalamic activity G: Two example PSTHs showing activated (left) and inhibited (right) thalamic neurons H: The mean z-scored firing for all activated and inhibited thalamic neurons.

**Figure S4:**
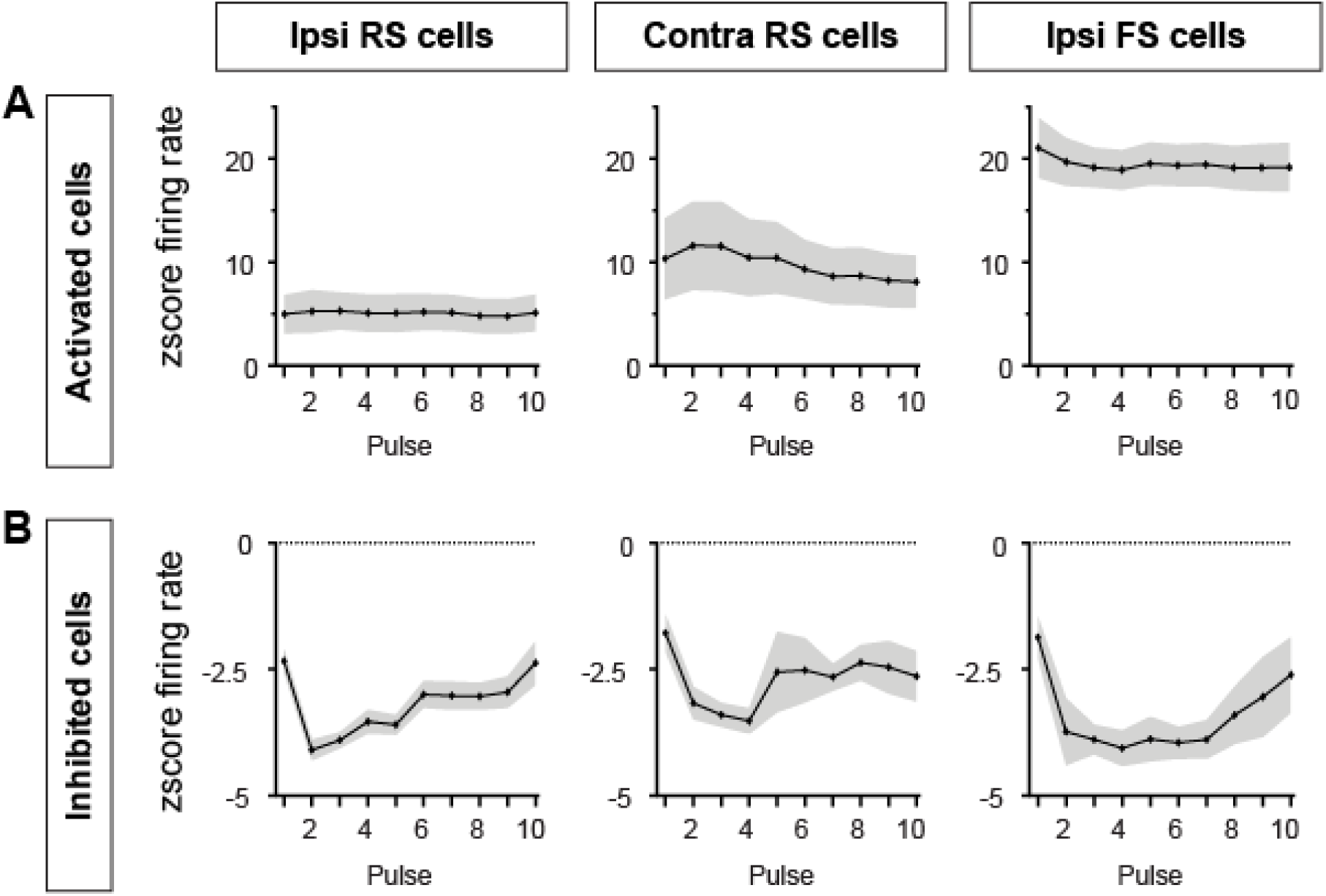
Detailed analysis of the claustrum RS cells shown in Figure 2 and FS cells shown in Figure 3 in terms of their response to ACC inputs. A: Activated cells in the contralateral hemisphere are more entrained to the ACC stimulation and show a trend toward being more positively modulated. B: Inhibited cells in the ipsilateral RS are more negatively modulated (more suppressed) and are less well entrained to the ACC stimulation showing that the effective feed-forward inhibition in the contralateral claustrum is weaker than ipsilateral claustrum.

**Figure S5:**
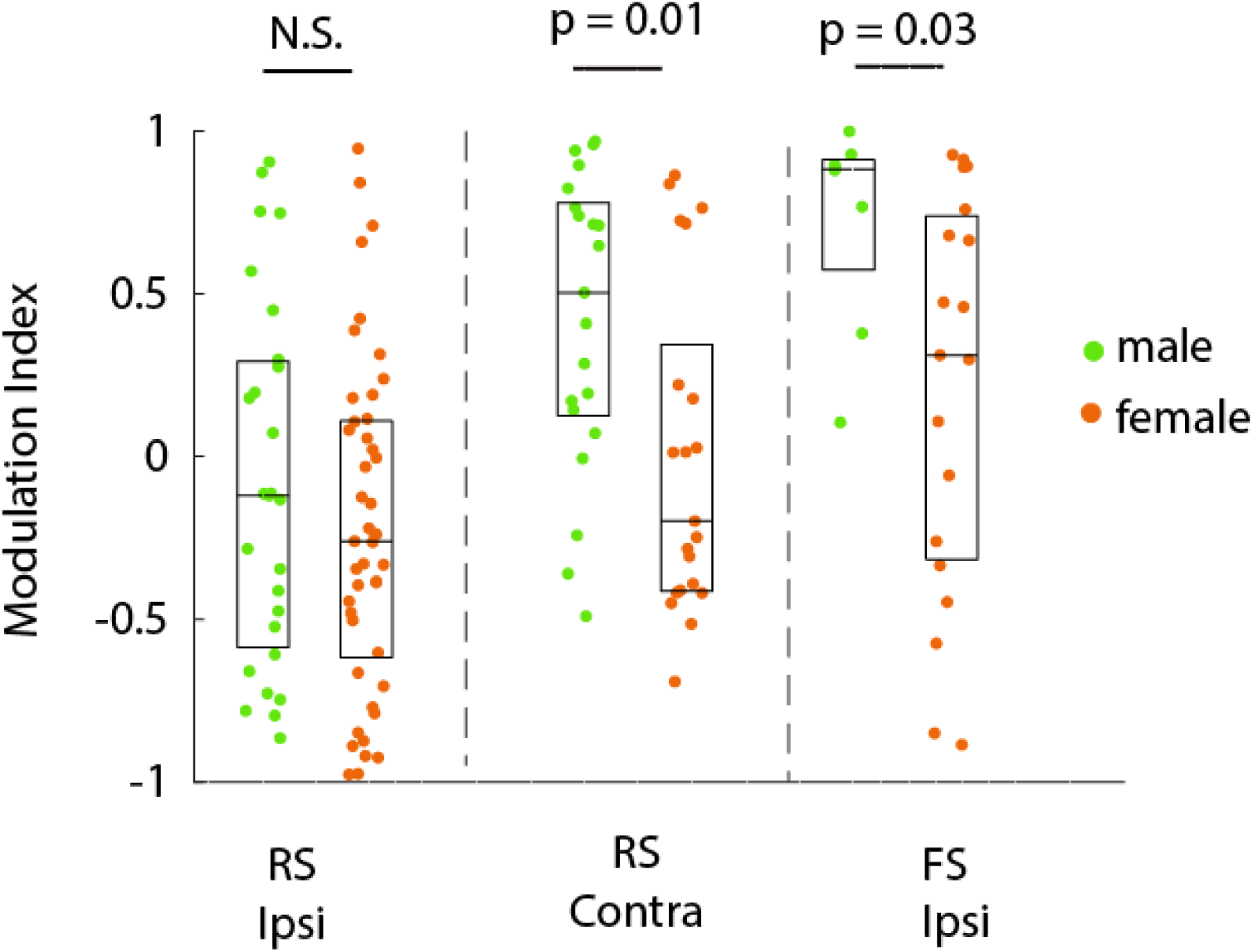
Analysis of how RS and FS cells respond to ACC stimulation as a function of sex. Modulation index of claustrum neurons during ACC stimulation is separately plotted for male and female adult mice. In the main text/figures, neurons for both sexes are merged.

**Figure S6:**
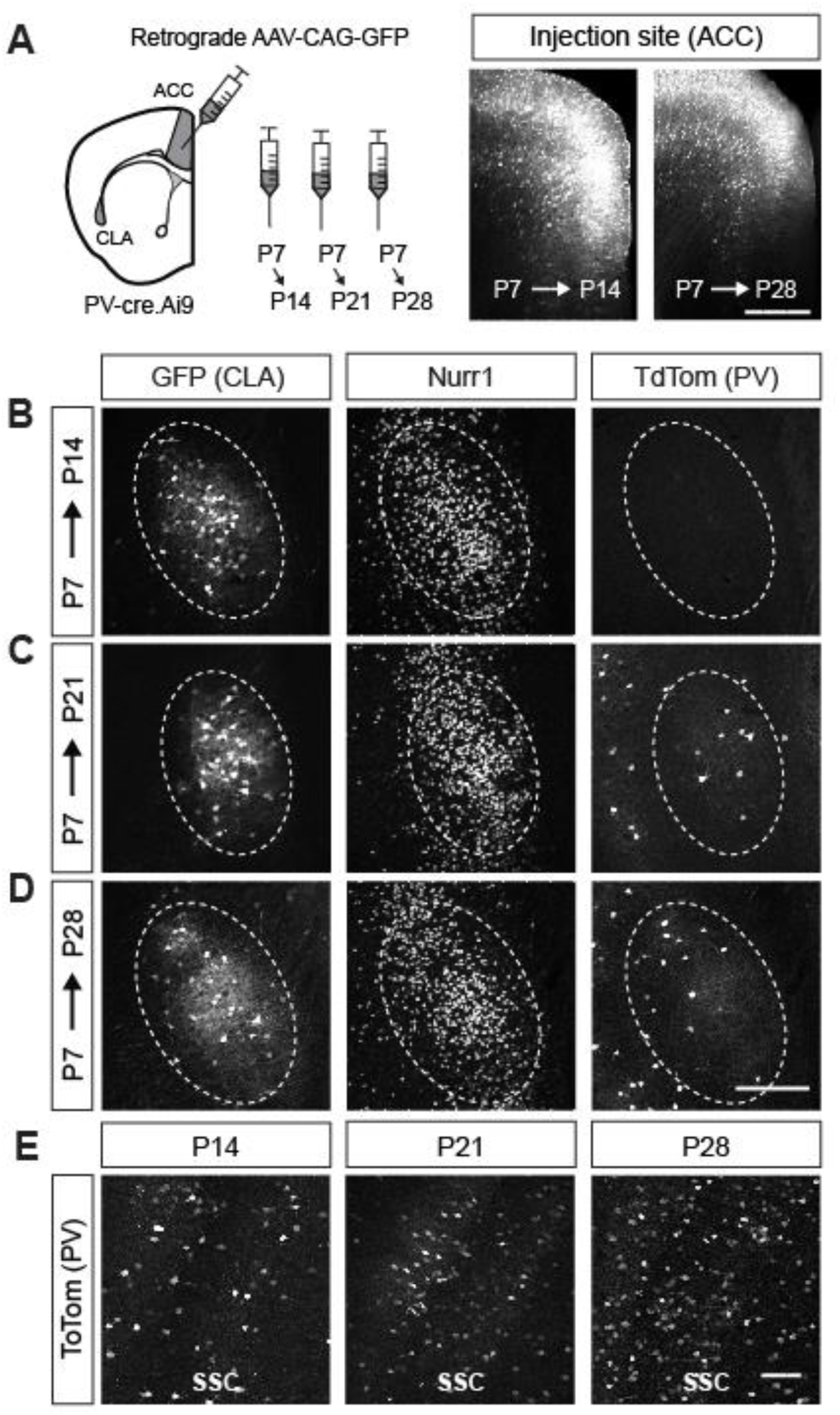
Developmental characterization of parvalbumin expressing cells in the claustrum and cortex of PV-cre:Ai14 mouse line. A: Schematic of retrograde tracer injection into the ACC at different ages to demarcate the claustrum (CLA). Mice were perfused 7, 14 and 21 days later. Two examples of virus injection sites are shown in different mice at P14 and P28 following injection. B-D: Retrograde tracer (GFP) and Nurr1 expression help identify the claustrum across age. Note the lack of PV at P14 in B. E: PV cells were identified in the somatosensory (SSC) cortex, just dorsal to the claustrum, at all ages. Scale bar in A: 400um, in B-D: 250um, E: 200um.

**Figure S7:**
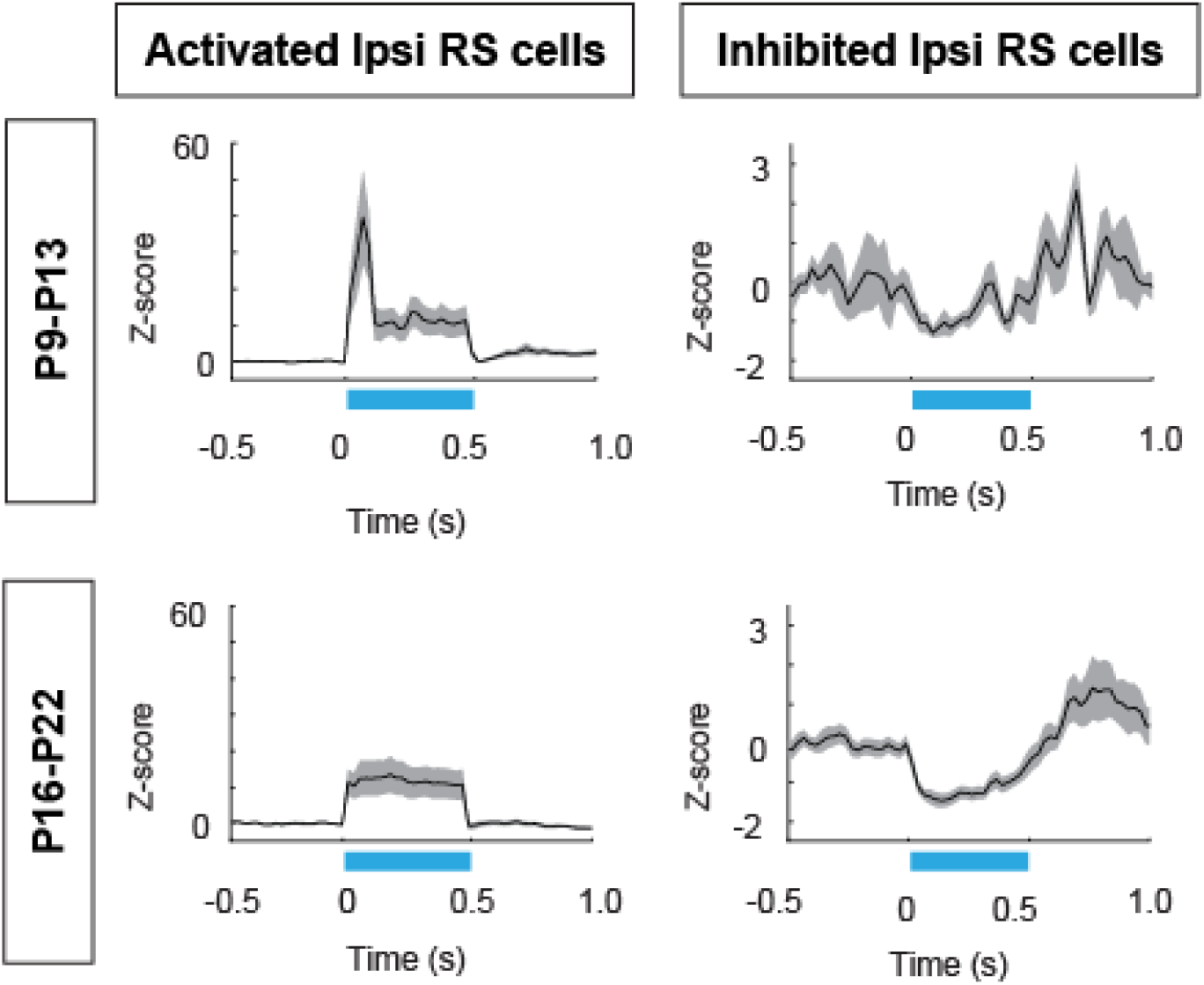
The mean z-scored firing rate of modulated cells in the claustrum as a function of age during postnatal development. Activated (left) and inhibited (right) neurons were sorted at distinct ages based on the state of PV cell maturation (immature PV cells: P9-P13; maturing PV cells: P16-P22). Note the strong activation of ipsilateral claustrum RS cells during the stimulation onset that is lost as juvenile mice begin the adolescence stage. Activated ipsi RS cells, P9-P13: n = 19, P16-P22: n = 9. Inhibited ipsi RS cells, P9-P13: n = 3, P16-P22: n = 21.

## Notes

### Competing Interest Statement

The authors have declared no competing interest.

